# SLC19A1 is an importer of the immunotransmitter cGAMP

**DOI:** 10.1101/539247

**Authors:** Anthony F. Cordova, Christopher Ritchie, Gaelen T. Hess, Michael C. Bassik, Lingyin Li

## Abstract

2’3’-cyclic-GMP-AMP (cGAMP) is a second messenger that activates the antiviral Stimulator of Interferon Genes (STING) pathway. We recently identified a novel role for cGAMP as a soluble, extracellular immunotransmitter that is produced and secreted by cancer cells. Secreted cGAMP is then sensed by host cells, eliciting an antitumoral immune response. Due to the antitumoral effects of cGAMP, other CDN-based STING agonists are currently under investigation in clinical trials for metastatic solid tumors. However, it is unknown how cGAMP and other CDNs cross the cell membrane to activate intracellular STING. Using a genome-wide CRISPR screen we identified SLC19A1 as the first known importer of cGAMP and other CDNs, including the investigational new drug 2′3′-bisphosphosphothioate-cyclic-di-AMP (2′3′-CDA^S^). These discoveries will provide insight into cGAMP’s role as an immunotransmitter and aid in the development of more targeted CDN-based cancer therapeutics.

## Introduction

Harnessing innate immunity to treat cancer is at the cutting edge of precise and personalized cancer treatment, and there is mounting evidence that the cGAMP-STING innate immunity pathway is a potent anti-cancer target (Wang et al. 2017; Deng et al. 2014; Corrales et al. 2015). The cyclic dinucleotide (CDN) cGAMP is a second messenger that is synthesized by cyclic-GMP-AMP synthase (cGAS) after detection of double-stranded DNA (dsDNA) in the cytosol (Sun et al. 2013). cGAMP binds and activates the cytosolic domain of its ER-membrane receptor STING, which in turn activates TBK1, a kinase, and IRF3, a transcription factor, resulting in the transcription, expression, and secretion of cytokines such as interferon-beta (IFN-β). These potent antiviral and anticancer cytokines can directly neutralize threats (Apelbaum et al. 2013) and trigger downstream adaptive immunity (Iwasaki and Medzhitov 2010). In the case of cancer clearance, IFN-β promotes cross-priming of CD8^+^ T cells by tumor-infiltrating antigen presenting cells (Fuertes et al. 2011). Primed CD8^+^ T cells can then infiltrate and kill both primary and metastatic tumors, leading to systemic tumor regression and long-term humoral memory of the tumor (Corrales et al. 2015; Woo et al. 2014).

While cytosolic dsDNA was originally discovered as a signal of viral infection (X.-D. Li et al. 2013), it is now also recognized as a hallmark of cancer (Mackenzie et al. 2017; Bakhoum et al. 2018). Cancer cells often have unstable genomes that result in improper chromosome segregation during mitosis. This leads to the formation of micronuclei enclosed by leaky membranes, thereby exposing dsDNA to the cytosol and activating the cGAMP-STING pathway (Mackenzie et al. 2017; Harding et al. 2017). Instead of inactivating the pathway to escape immune detection, the vast majority of cancer cells retain the STING pathway (Bakhoum and Cantley 2018) and exploit it to their advantage in at least two ways. First, cGAS promotes cancer progression by inhibiting DNA repair (Liu et al. 2018), thereby increasing genomic instability. Second, many cancer cells rewire the STING pathway to promote metastasis, while avoiding IFN-β production (Bakhoum and Cantley 2018; Bakhoum et al. 2018).

While cancer cells do not typically produce type I interferons, it has been shown that cGAMP-producing cancer cells can activate the STING pathway in nearby *cGAS*^-/-^ host cells (Marcus et al. 2018). Although the mechanism for this was previously unknown, we recently determined that it is due to cGAMP’s role as an immunotransmitter between cancer and host cells (Carozza et al., in revision). Cancer cells are able to efficiently export cGAMP into the extracellular space, which can then be degraded by the extracellular hydrolase ENPP1 (L. Li et al. 2014), or can cross the cell membrane of nearby host cells through unknown mechanisms, activating STING and the downstream anti-cancer immune response.

As cGAMP can elicit a robust antitumoral immune response, there has been great interest in developing cGAMP analogs as cancer therapeutics. In 2014, we reported stable phosphothioate analogs of cGAMP (L. Li et al. 2014). Subsequent studies have shown that intratumoral injections of these and other phosphothioate analogs can cure multiple tumor types in mice (Corrales et al. 2015). With these promising results, 2’3’-bisphosphothioate-cyclic-di-AMP (2’3’-CDA^S^) and another stable CDN analog entered phase-I clinical trials in 2017, both in combination with PD-1 checkpoint inhibitors (Trial IDs NCT03172936 and NCT03010176, respectively). Despite their therapeutic potential, it is still unknown how cGAMP and other CDNs enter the cytoplasm of target cells. Because they are unable to passively diffuse across the plasma membrane due to their negative charges, they must enter cells through a facilitated mechanism. This mechanism could be either specific (e.g. through a transporter) or non-specific (e.g. pinocytosis).

Understanding how cGAMP enters cells is essential for both characterizing cGAMP as an immunotransmitter and developing CDN-based STING agonists as therapeutics. Here, we describe SLC19A1 as the first known transporter of the immunotransmitter cGAMP. In addition, we show that SLC19A1 is also an importer of several bacterial and synthetic CDNs.

## Results

### A genetic screen identifies putative components of the extracellular cGAMP-STING pathway

To identify important factors for cGAMP internalization, we performed a whole-genome CRISPR screen in the monocyte-derived U937 cell line. U937 cells were chosen for this screen because monocyte-lineage cells have high levels of STING and a higher response to exogenous STING agonists *in vivo* compared to other immune cells (Sivick et al. 2018). U937 cells express all STING pathway components (**Supplementary Fig. 1a**) and respond to extracellular cGAMP by phosphorylating the transcription factor IRF3 (**Fig. 1a**) and producing IFN-β (**Supplementary Fig. 1b-1c**). Importantly, the response is independent of the cGAMP synthase cGAS, suggesting that it is due to exogenous, extracellular cGAMP (**Supplementary Fig. 1d**). With prolonged cGAMP treatment we found that U937 cells die in a dose-dependent manner, making them well-suited for a live/dead CRISPR screen (**Fig. 1b**).

**Figure 1.**
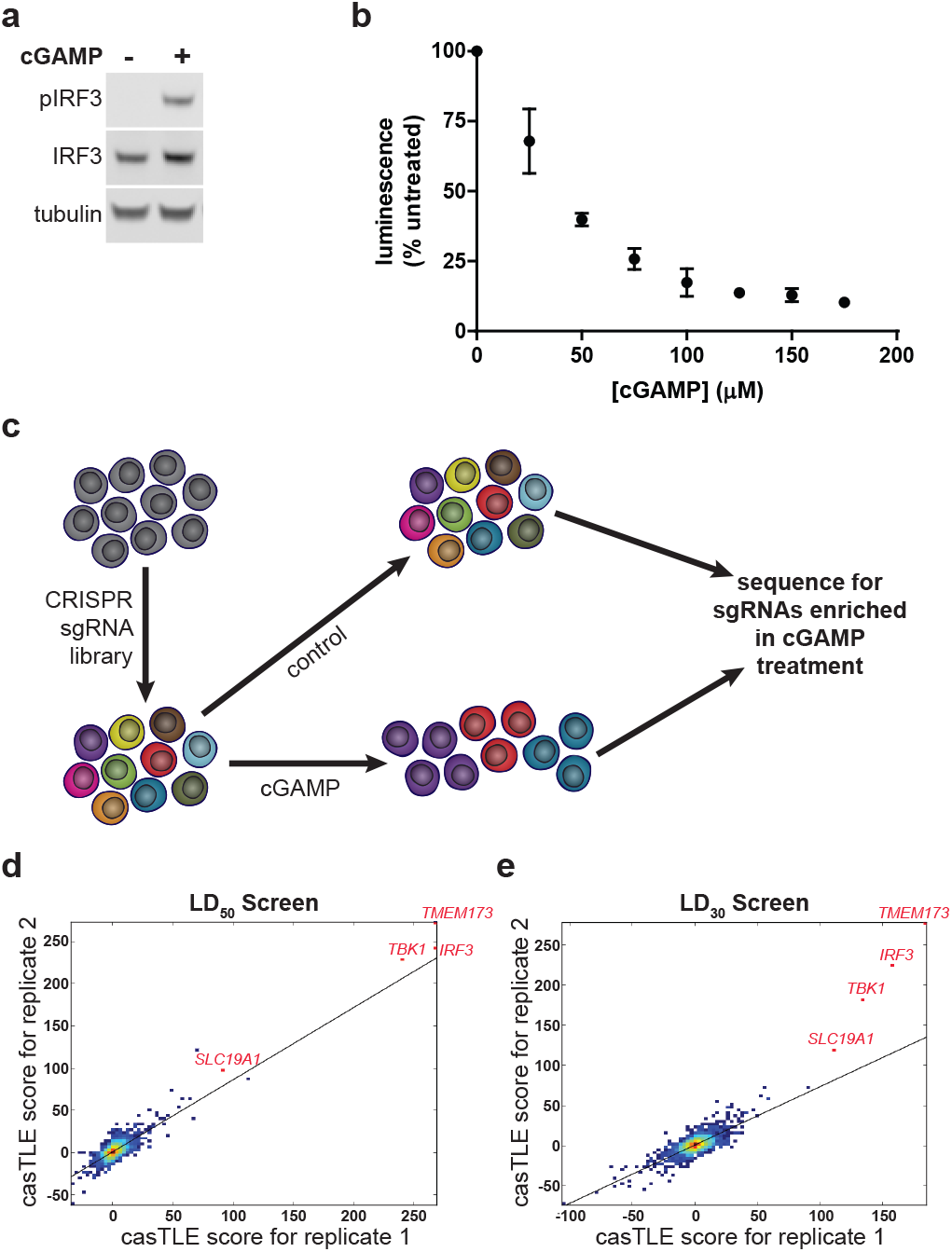
A genetic screen identifies putative components of the extracellular cGAMP-STING pathway. (**a)** IRF3 phosphorylation in response to extracellular cGAMP. U937 cells were treated with 100 μM cGAMP for 2 h. (**b)** U937 cells were treated with various concentrations of cGAMP for 16 h. Cell viability was assessed using CellTiter-Glo (n = 2 biological replicates). (**c)** Schematic of the CRISPR screen. A whole-genome sgRNA library was created in U937 cells. 250 million library cells were treated with cGAMP every 24 h for 2 weeks. The genomic DNA was sequenced and highly-enriched sgRNA sequences were identified. (**d-e)** casTLE scores of individual enriched genes from (**d)** two replicates using LD_50_ doses of cGAMP and (**e)** two replicates using LD_30_ doses of cGAMP. Top hits are annotated in red.

In previous work, we had successfully designed and introduced a custom genome-wide single-guide RNA (sgRNA) library into U937 cells to identify genes required for cell growth (Morgens et al. 2017). Additionally, we were able to synthesize hundreds of milligrams of high purity cGAMP using previously developed methods (L. Li et al. 2014; Civril et al. 2013). This enabled us to perform a live/dead screen by treating the library cells with a cGAMP concentration sufficient to kill 50% of cells (LD_50_) every 24 hours for 2 weeks. In this screen, cells containing sgRNAs that target key STING pathway members, including cGAMP importers, should be resistant to cGAMP treatment and enriched (**Fig. 1c**). Therefore, by sequencing the sgRNAs in treated and untreated populations to calculate relative enrichment and depletion of sgRNAs, we generated a list of candidate genes involved in the extracellular cGAMP-STING pathway. We also repeated the screen with a lower cGAMP concentration (LD_30_) and obtained similar results. The casTLE score (a measure of statistical significance for the collective enrichment of sgRNAs targeting a particular gene (Morgens et al. 2016)) was reproducible between replicates (**Fig 1d-e**). The top three enriched genes in both screens were the known STING pathway members *TMEM173, TBK1*, and *IRF3*, further supporting the validity of this screening method (**Supplementary Fig. 1e**).

### SLC19A1 is essential for robust extracellular cGAMP signaling in U937 cells

Among the top hits from our CRISPR screen, we focused on SLC19A1 as a candidate transporter of cGAMP. The SLC19A1 protein, also known as Reduced Folate Carrier 1 (RFC1), has been previously characterized as a high affinity importer of reduced folates, such as folinic acid, and antifolates, such as methotrexate (Hou and Matherly 2014; G Jansen et al. 1990). Given the structural similarity between cGAMP and these known SLC19A1 substrates (**Fig. 2a**), we hypothesized that SLC19A1 also imports cGAMP.

**Figure 2.**
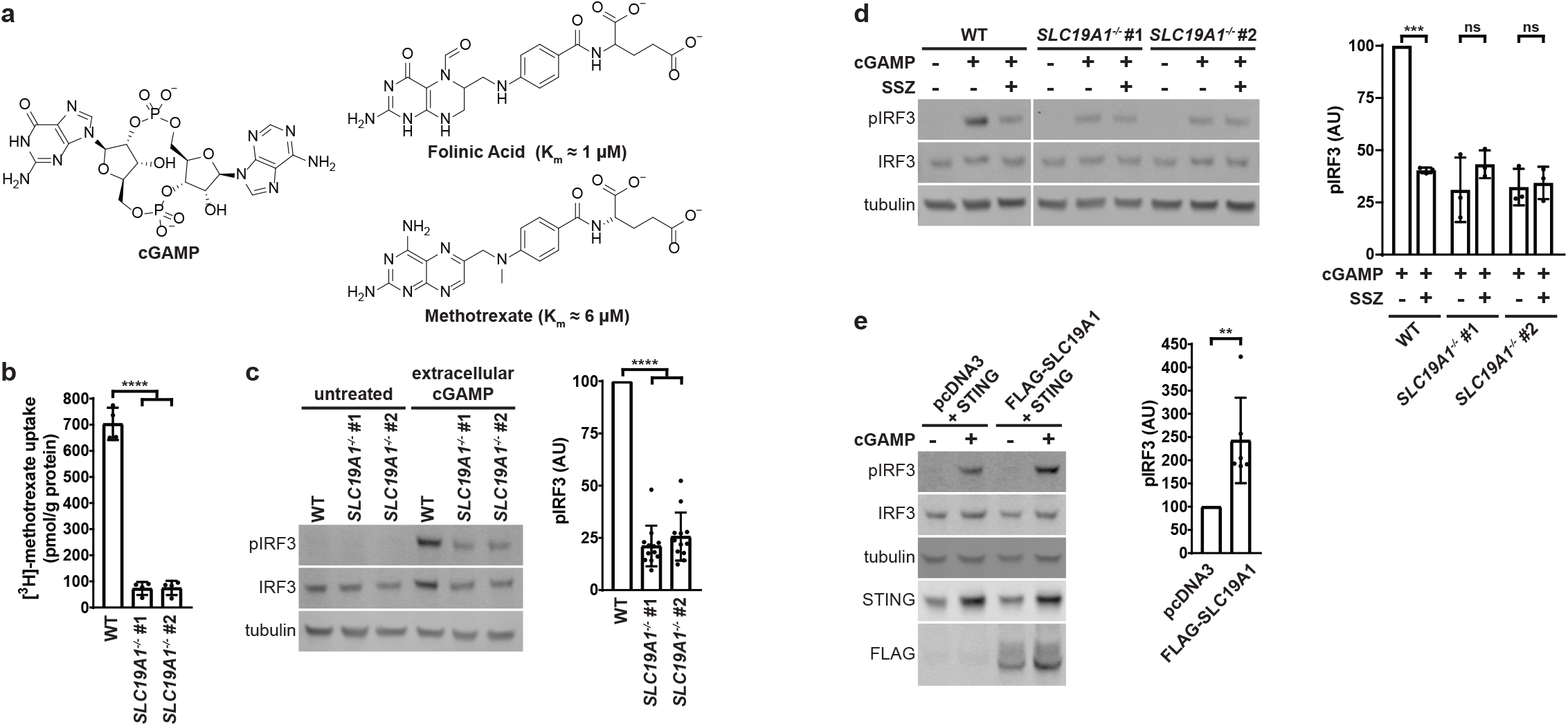
SLC19A1 is essential for robust extracellular cGAMP signaling in U937 cells. (**a)** Chemical structures of the known SLC19A1 substrates folinic acid and methotrexate, as compared to cGAMP. (**b)** Uptake of [^3^H]-methotrexate in U937 *SLC19A1*^-/-^ lines. U937 WT and *SLC19A1*^-/-^ lines were treated with 17.5 nM [^3^H]-methotrexate for 5 min. Results are shown as the mean ± SD (n = 4 biological replicates). (**c)** Effect of *SLC19A1* knockout on extracellular cGAMP signaling. U937 WT and *SLC19A1*^-/-^ cells were treated with 100 μM cGAMP for 2 h. Representative Western blot is shown with quantification of mean ± SD (n = 12 biological replicates). (**d)** Effect of sulfasalazine (SSZ) on extracellular cGAMP signaling. WT and *SLC19A1*^-/-^ U937 cells were pretreated with 1 mM SSZ for 20 min followed by a 100 μM cGAMP treatment for 2 h. Representative Western blot is shown with quantification of mean ± SD (n = 3 biological replicates). (**e)** Effect of SLC19A1 overexpression on extracellular cGAMP signaling. HEK 293T cells were transfected with pcDNA3-STING-HA and either an empty pcDNA3-FLAG-HA vector or pcDNA3-FLAG-HA-SLC19A1. Transfected cells were treated 24 h after transfection with 100 μM cGAMP for 2 h. Representative Western blot is shown with quantification of mean ± SD (n = 6 biological replicates).

We first sought to determine if SLC19A1 is important for extracellular cGAMP signaling by using CRISPR/Cas9 to knock out *SLC19A1* in U937 cells. Because there are no commercial antibodies that effectively detect SLC19A1, we sequenced the *SLC19A1* locus and identified two *SLC19A1*^-/-^ clones containing frameshift mutations that lead to early termination (**Supplementary Fig. 2a-b**). In contrast to the parent U937 cell line, these *SLC19A1*^-/-^ clones no longer import [^3^H]-methotrexate, demonstrating the loss of functional SLC19A1 protein (**Fig. 2b**). Importantly, protein levels of known essential STING pathway members were not significantly altered in these *SLC19A1*^-/-^ clones (**Supplementary Fig. 2c**). When treated with extracellular cGAMP, *SLC19A1*^-/-^ cells showed 75% less IRF3 phosphorylation than wild type cells (**Fig. 2c**), as well as greatly diminished production of IFN-β mRNA (**Supplementary Fig. 2d**) and protein (**Supplementary Fig. 2e**). In addition, the non-competitive SLC19A1 inhibitor sulfasalazine (Gerrit Jansen et al. 2004) reduced phosphorylation of IRF3 in response to extracellular cGAMP by 60% in wild type U937 cells, but not in *SLC19A1*^-/-^ cells (**Fig. 2d**). Our genetic and pharmacological data together support an important role for SLC19A1 in extracellular cGAMP signaling.

To determine if SLC19A1 is sufficient to increase extracellular cGAMP signaling, HEK 293T cells (which lack endogenous cGAS and STING, and are therefore more amenable to transfection than U937 cells) were co-transfected with pcDNA3-STING-HA and either an empty pcDNA3-FLAG-HA vector or pcDNA3-FLAG-HA-SLC19A1. Overexpression of SLC19A1 increased the response to extracellular cGAMP by over 200% compared to transfection with the empty vector (**Fig. 2e**), indicating that SLC19A1 expression is sufficient to increase extracellular cGAMP signaling.

### SLC19A1 facilitates extracellular uptake of cGAMP

To determine whether SLC19A1 is required for extracellular cGAMP uptake or for downstream STING signaling, we examined its role in intracellular cGAMP signaling. While direct detection of cGAMP import would be ideal, the current technologies to measure cGAMP concentrations using ^32^P-radiolabeled CDNs (L. Li et al. 2014) or mass spectrometry (Carozza et al., in revision) are not sensitive enough to detect nanomolar concentrations of intracellular cGAMP, given the high background cGAMP in the media. However, STING binds to cGAMP with a *K*_d_ of ∼5 nM (Zhang et al. 2013). Therefore, the best readout of cGAMP internalization is still STING activation (Ablasser et al. 2013; Bridgeman et al. 2015), as quantified by IRF3 phosphorylation. Intracellular cGAMP was introduced into U937 cells through cGAMP electroporation (**Fig. 3a**). In contrast to their response to extracellular cGAMP, U937 *SLC19A1*^-/-^ cells are not defective in intracellular cGAMP signaling, as evidenced by their response to electroporated cGAMP (**Fig. 3b**). These results indicate that SLC19A1 facilitates uptake of extracellular cGAMP.

**Figure 3.**
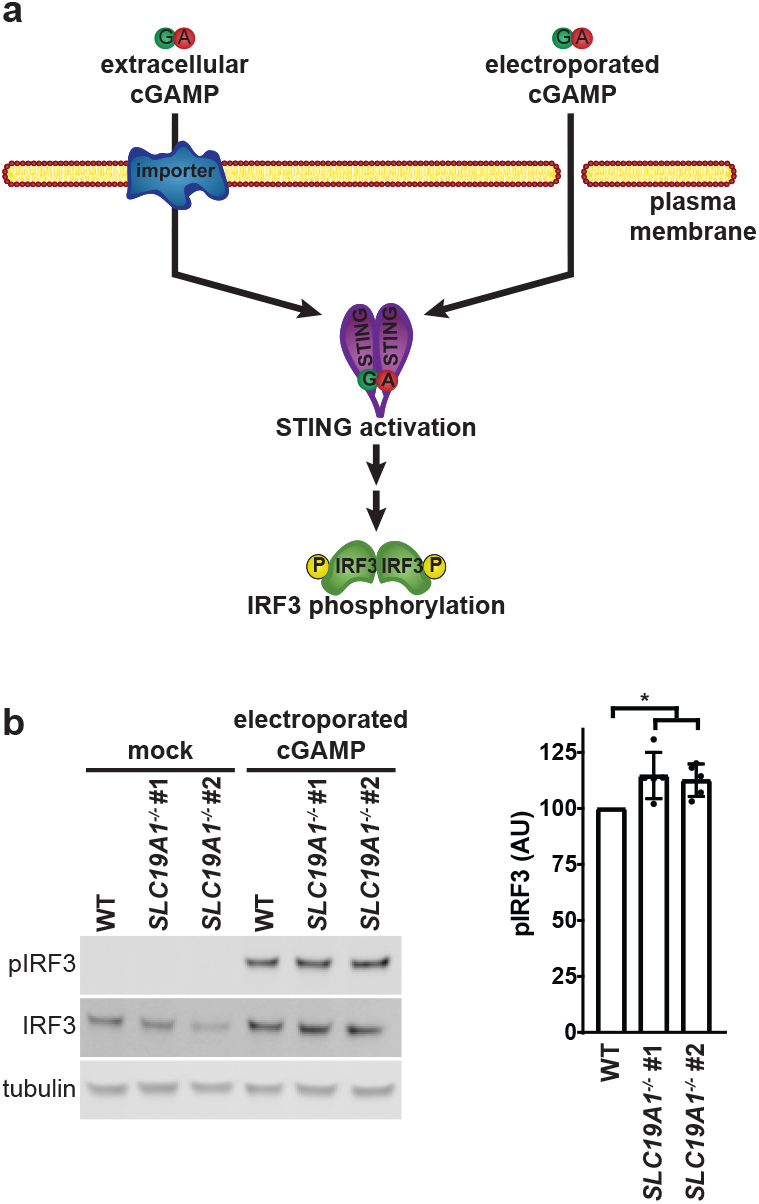
SLC19A1 facilitates extracellular uptake of cGAMP. (**a)** Schematic of bypassing cGAMP import through cGAMP electroporation. (**b)** U937 WT and *SLC19A1*^-/-^ cells were electroporated with 100 nM cGAMP for 2 h. Representative Western blot is shown with quantification of mean ± SD (n = 5 biological replicates).

### SLC19A1 is a direct cGAMP importer

We then tested whether SLC19A1 is a direct cGAMP importer requiring the transporter activity of the protein, rather than a receptor for endocytosis or a scaffold for the direct importer. SLC19A1 is an antiporter, and export of intracellular organic anions facilitates uptake of its substrates (Hou and Matherly 2014). Through this mechanism, the cell-permeable compound 5-amino-4-imidazolecarboxamide riboside (AICAR) boosts import of SLC19A1 substrates. Once inside the cell, AICAR is phosphorylated to AICAR monophosphate, which serves as an organic anion for SLC19A1’s antiporter activity (Visentin, Zhao, and Goldman 2012). Treating U937 cells with AICAR increased the response to extracellular cGAMP by approximately 200% in an SLC19A1-dependent manner (**Fig. 4a**). While AICAR is known to cause AMPK phosphorylation in some cells (Sullivan et al. 1994), which may affect STING signaling through a negative feedback loop (Konno, Konno, and Barber 2013), it did not have any additional effect on AMPK phosphorylation in cGAMP-treated U937 cells (**Supplementary Fig. 4**).

**Figure 4.**
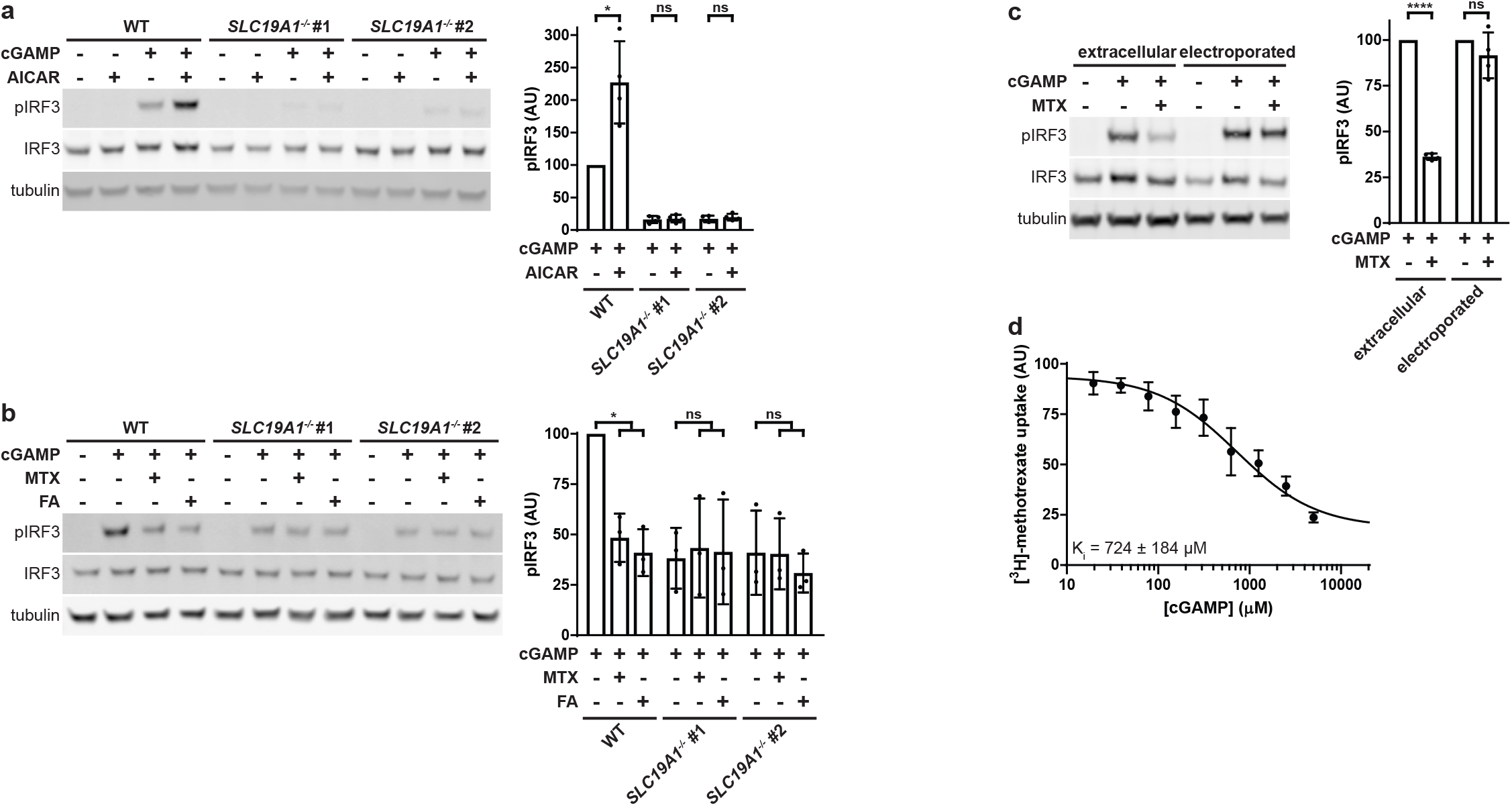
SLC19A1 is a direct cGAMP importer. (**a)** Effect of AICAR on extracellular cGAMP signaling. U937 WT and *SLC19A1*^-/-^ cells were pretreated with 1 mM AICAR for 20 min followed by treatment with 100 μM 2′3′-cGAMP for 2 h. Representative Western blot is shown with quantification of mean ± SD (n = 4 biological replicates). (**b)** Effect of competitive SLC19A1 inhibitors on extracellular cGAMP signaling. U937 WT and *SLC19A1*^-/-^ cells were treated with 100 μM cGAMP alone or in the presence of 500 μM methotrexate (MTX) or 500 μM folinic acid (FA) for 2 h. Representative Western blot is shown with quantification of mean ± SD (n = 3 biological replicates). (**c)** Bypass of MTX inhibition by electroporation. U937 WT cells were pretreated with 500 μM MTX for 5 min., then treated with either 100 μM extracellular cGAMP or electroporated with 100 nM cGAMP for 90 min. Representative Western blot is shown with quantification of mean ± SD (n=4). (**d)** Effect of cGAMP on [^3^H]-methotrexate uptake. U937 cells were treated 17.5 nM [^3^H]-methotrexate in the presence of multiple concentrations of cGAMP for 1 min (n = 3 biological replicates).

Next, we tested whether the well-characterized SLC19A1 substrates folinic acid and methotrexate could act as competitive inhibitors of cGAMP to reduce extracellular cGAMP signaling. Although folinic acid and methotrexate have opposing roles in folate metabolism, they both reduced extracellular cGAMP-induced IRF3 phosphorylation by more than 50% in an SLC19A1-dependent manner (**Fig. 4b**). In addition, methotrexate had no effect on IRF3 phosphorylation when cGAMP was electroporated into cells, indicating that methotrexate inhibits cGAMP import (**Fig. 4c**). Furthermore, cGAMP inhibited import of [^3^H]-methotrexate, with a K_i_ of ∼724 μM (**Fig. 4d**). Together, our results demonstrate that SLC19A1 is a direct cGAMP importer and the dominant one in U937 cells. However, the presence of residual IRF3 phosphorylation in extracellular cGAMP-treated *SLC19A1*^-/-^ cells suggests that there are one or more additional cGAMP import mechanisms in U937 cells.

### SLC19A1 imports bacterial and synthetic CDNs, including 2’3’-CDA^S^

In addition to cGAMP, bacterial CDN second messengers, including 3′3′-cyclic-GMP-AMP (3′3′-cGAMP) (Zhang et al. 2013), 3′3′-cyclic-di-GMP (3′3′-CDG) (Burdette et al. 2011), and 3′3′-cyclic-di-AMP (3′3′-CDA) (Woodward, Iavarone, and Portnoy 2010), also bind and activate the STING pathway (**Fig. 5a**). These CDNs are critical for the immune detection of pathogens and contribute to pathogen virulence in mice (Sauer et al. 2011; Dey et al. 2015; Woodward, Iavarone, and Portnoy 2010; Davies et al. 2012). It is unclear how these CDNs are exposed to STING in the cytosol since most bacterial pathogens, such as the 3′3′-cGAMP-producing *Vibrio cholera*, are extracellular, and many intracellular bacteria, such as the 3’3’-CDA-producing *Mycobacterium tuberculosis*, hide in intracellular compartments to avoid detection. Given the structural similarities shared by these bacterial CDNs and cGAMP, we hypothesized that they are also SLC19A1 substrates. As expected, extracellular 3’3’-cGAMP signaling was diminished by approximately 70% in *SLC19A1*^-/-^ cells relative to wild type, indicating that 3’3’-cGAMP is an SLC19A1 substrate (**Fig. 5b**). However, U937 cells responded poorly to extracellular 3’3’-CDA and 3’3’-CDG compared to other CDNs (**Supplementary Fig. 5a**), limiting our ability to assess their dependence on SLC19A1 for import.

**Figure 5.**
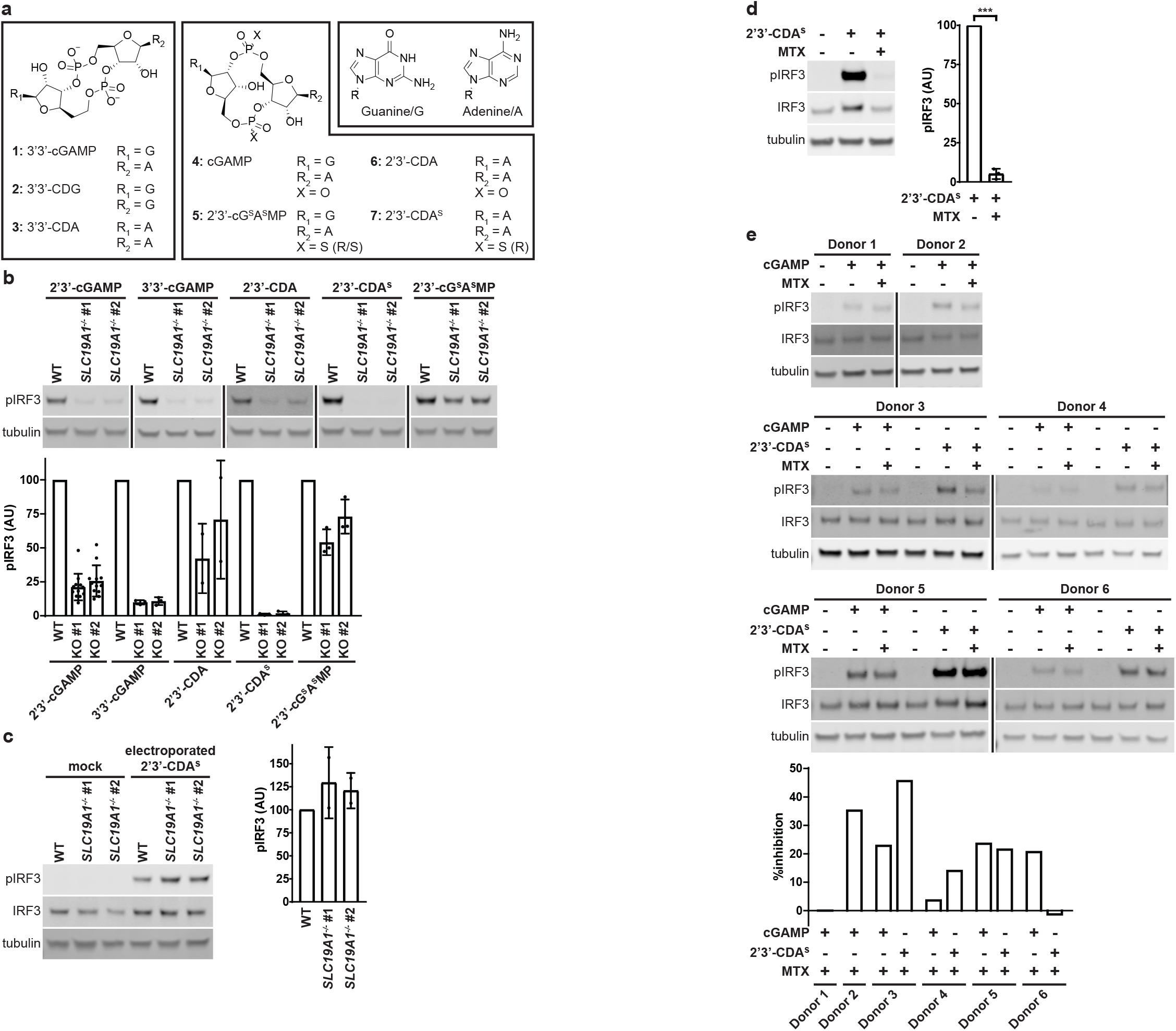
SLC19A1 imports bacterial and synthetic CDNs, including 2’3’-CDA^S^. (**a)** Structures of bacterial (1-3), mammalian (4), and synthetic (5-7) CDNs. (**b)** Importance of SLC19A1 for signaling of various extracellular CDNs. U937 WT and *SLC19A1*^-/-^ cells were treated with 200 μM 3′3′-cGAMP (n = 3 biological replicates) for 3 h, or with either 100 μM cGAMP (n = 12 biological replicates), 25 μM 2′3′-cG^S^A^S^MP (n = 3 biological replicates), 100 μM 2’3’-CDA (n = 2 biological replicates), or 15 μM 2’3’-CDA^S^ (n = 3 biological replicates) for 2 h. Representative Western blots are shown with quantification of means ± SD. (**c)** Bypass of 2’3’-CDA^S^ import by electroporation. U937 WT and *SLC19A1*^-/-^ cells were electroporated with 100 nM 2’3’-CDA^S^ for 2 h. Representative Western blot is shown with quantification of mean ± SD (n = 2 biological replicates). (**d)** Effect of MTX on extracellular 2’3’-CDA^S^ signaling. U937 cells were treated with 500 μM MTX and 15 μM 2’3’-CDA^S^ for 90 min. Representative Western blot is shown with quantification of mean ± SD (n = 3 biological replicates). (**e)** Effect of MTX on extracellular cGAMP signaling in primary CD14^+^ monocytes. CD14^+^ monocytes were isolated from six different healthy donors. Monocytes were pretreated with 500 μM MTX for 5 min then treated with either 50 μM cGAMP or 12.5 μM 2’3’-CDA^S^ for 60 min. Response to 2’3’-CDA^S^ was not tested in Donors 1 and 2. Quantifications are shown as percent inhibition of pIRF3 signal with MTX pretreatment relative to no pretreatment.

We then tested whether SLC19A1 transports synthetic cGAMP analogs, including 2′3′-bis-phosphothioate-cyclic-GMP-AMP (2′3′-cG^S^A^S^MP) (L. Li et al. 2014), 2′3′-cyclic-di-AMP (2’3’-CDA), and the investigational new drug 2’3’-CDA^S^ (**Fig. 5a**). All of these 2’3’-CDNs elicited a weaker response in *SLC19A1*^-/-^ cells than in wild type U937 cells, indicating that they are SLC19A1 substrates (**Fig. 5b**). Interestingly, these substrates have different selectivities for SLC19A1, with 2’3’-CDA^S^ being the most selective, followed by cGAMP, and 2’3’-CDA and 2′3′-G^S^A^S^MP being the least selective. Strikingly, loss of SLC19A1 completely abolished STING signaling in response to extracellular 2’3’-CDA^S^ but did not reduce the response to electroporated 2’3’-CDA^S^ (**Fig. 5c**). Consistent with this finding, extracellular 2’3’-CDA^S^ signaling was also greatly reduced in the presence of methotrexate, demonstrating that SLC19A1 is necessary for 2’3’-CDA^S^ import in U937 cells (**Fig. 5d, Supplementary Fig. 5b**). To determine if SLC19A1 is sufficient for 2’3’-CDA^S^ import, we co-transfected HEK 293T cells with a STING plasmid and either FLAG-HA-SLC19A1 or an empty vector. While cells transfected with an empty vector responded poorly to extracellular 2’3’-CDA^S^, cells overexpressing SLC19A1 had a robust response (**Supplementary Fig. 5c**).

Interestingly, electroporation of an equimolar amounts of cGAMP and 2’3’-CDA^S^ resulted in significantly less IRF3 phosphorylation in cells electroporated with 2’3’-CDA^S^, as compared to cGAMP. In contrast, treatment with 15 μM extracellular 2’3’-CDA^S^ resulted in significantly higher levels of phosphorylated IRF3 than treatment with 100 μM extracellular cGAMP (**Supplementary Fig. 5d**). This suggests that the increased response to extracellular 2’3’-CDA^S^ is due to it being a higher-affinity substrate for SLC19A1 than cGAMP, rather than due to stronger STING activation once inside cells.

### SLC19A1 is involved in extracellular CDN signaling in primary CD14^**+**^ cells

Because SLC19A1 is the dominant cGAMP importer in monocyte-derived U937 cells, we sought to characterize its role in primary human CD14^+^ monocytes. As attempts to knockdown SLC19A1 using siRNA and CRISPR/Cas9 proved to be toxic to monocytes, we relied on pharmacological inhibition of SLC19A1 with methotrexate. CD14^+^ monocytes were isolated from the whole blood of six individual donors. The inhibitory effect of methotrexate on the response to extracellular cGAMP and 2’3’-CDA^S^ varied between donors, ranging from 0% inhibition to 35% (for cGAMP) and 45% (for 2’3’-CDA^S^) inhibition (**Fig. 5e**). These data suggest that SLC19A1 also plays a role in CDN import in primary monocytes, and that its use varies among donors.

## Discussion

In this study we performed a whole-genome CRISPR screen in the monocyte-derived U937 cell line to identify key regulators of the response to extracellular cGAMP. From the screen we identified SLC19A1 as a cGAMP importer, providing the first evidence of a specific import mechanism for cGAMP. In addition to importing cGAMP, we showed that SLC19A1 also imports other CDNs, such as the investigational new drug 2’3’-CDA^S^. We found that the selectivity of CDNs for SLC19A1 is dependent on the identity of nucleotide bases and linkages. For example, whereas 3’3’-cGAMP is selective for SLC19A1, the other 3’3’-linked CDNs, 3’3’-CDA and 3’3’-CDG, do not appear to be imported by SLC19A1. Similarly, while the phosphodiester-linked 2’3’-CDA is only moderately selective for SLC19A1 in U937 cells, the phosphothioate linked 2’3’-CDA^S^ is highly selective for SLC19A1 in this cell line.

Interestingly, different cell types use different sets of transport mechanisms. For example, while U937 cells primarily depend on SLC19A1 to import cGAMP, CD14^+^ monocytes use additional import mechanisms. Furthermore, different CDNs do not always enter the same cell type to the same degree. For example, while cGAMP can efficiently enter HEK 293T cells, 2’3’-CDA^S^ cannot. We therefore hypothesize that cGAMP utilizes a specific transporter in HEK 293T cells that has low affinity toward 2’3’-CDA^S^. Although the identity of the HEK 293T importer(s) remains unknown, it is unlikely that import is due to a nonspecific mechanism, such as pinocytosis, as this would not explain the differential response to extracellular cGAMP and 2’3’-CDA^S^.

In tumors, it has been shown that both endogenous and exogenous extracellular cGAMP promotes immune cell recruitment and tumor shrinkage in a STING-dependent manner (Corrales et al. 2015; Woo et al. 2014; Carozza et al., in revision). However, it still unknown which set of cGAMP-sensing cells are required for the antitumoral immune response, as well as if this set of cells varies by patient or tumor type. Future identification of additional transport mechanisms in different cell types will help uncover the identity of the responder cells.

The effects of cGAMP on tumor shrinkage in mice has led to the development of multiple CDN-based STING agonists. However, preliminary results from monotherapy CDN trials indicate that the drugs lack the robust antitumoral effect previously observed in mice (Meric-Bernstam et al. 2018; Harrington et al. 2018), and that different CDN analogs exhibited varying efficacies. One possible explanation is that these CDNs are unable to activate STING in responder cells due to inefficient cellular import. Since 2’3’-CDA^S^, the investigational new drug in one of these trials, is imported in part by SLC19A1, low SLC19A1 expression in responder cells could prevent 2’3’-CDA^S^ import and STING activation. Additionally, differential importer expression between patients may explain why some patients experienced a partial response (Meric-Bernstam et al. 2018), while others did not. As these importers are sensitive to subtle changes in CDN structures, it may be possible to design better CDN-based therapeutics that efficiently and selectively enter responder cells, leading to immune activation and tumor killing, while minimizing off-target adverse events.

## Materials and Methods

### Reagents and antibodies

2’3’-cyclic-GMP-AMP (cGAMP) was synthesized in house as described below. 3’3’-cyclic-GMP-AMP (3’3’-cGAMP), 3’3’-cyclic-di-AMP (3’3’-CDA), 3’3’-cyclic-di-GMP (3’3’-CDG), 2’3’-cyclic-di-AMP (2’3’-CDA), 2’3’-bisphosphothioate-cyclic-di-AMP (2’3’-CDA^S^), and 2’3’-bisphosphothioate-cyclic-GMP-AMP (2’3’-cG^S^A^S^MP) were purchased from Invivogen. 5-aminoimidazole-4-carboxamide ribonucleotide (AICAR), sulfasalazine, folinic acid, and methotrexate were purchased from Sigma Aldrich. Sulfasalazine was dissolved in 50 mM NaHCO_3_ and methotrexate was dissolved in 100 mM NaHCO_3_. CellTiter-Glo luminescent cell viability assay was purchased from Promega. Rabbit polyclonal antibodies against TBK1 (1:1000), IRF3 (1:1000), pTBK1 (S172, 1:1000), pIRF3 (S396, 1:1000), AMPKα (1:1000), pAMPKα (THr172, 1:1000), STING (1:1000), and cGAS (1:1000) were purchased from Cell Signaling Technology. Mouse monoclonal anti-α-tubulin (1:1000) was purchased from Cell Signaling Technology.

### Expression and purification of recombinant cGAS

The DNA sequence encoding porcine cGAS (residues 135-497) was amplified from a porcine cDNA library using the primer pair sscGAS FWD: 5’-CTGGAAGTTCTGTTCCAGGGGCCCCATATG GGCGCCTGGAAGCTCCAGAC-3’ and sscGAS REV: 5’-GATCTCAGTGGTGGTGGTGGTGGTGCTCGAG CCAAAAAACTGGAAATCCATTGT-3’. The PCR product was inserted into pDB-His-MBP (a generous gift from Qian Yin, Florida State University) via Gibson assembly and expressed in Rosetta cells. Cells were grown in 2xYT medium with 100 μg/mL kanamycin and induced with 0.5 mM IPTG when OD_600_ reached 1, and then were grown overnight at 16°C. All subsequent procedures using proteins and cell lysates were performed at 4°C. Cells were pelleted and lysed in 20 mM HEPES pH 7.5, 400 mM NaCl, 10% glycerol, 10 mM imidazole, 1 mM DTT, and protease inhibitors (cOmplete, EDTA-free protease inhibitor cocktail, Roche). Cell lysate was then cleared by ultracentrifugation at 50,000 x g for 1 h. The cleared supernatant was incubated with HisPur cobalt resin (ThermoFisher Scientific; 1 mL resin per 1 L bacterial culture) for 30 min. Cobalt resin was then washed with 20 mM HEPES pH 7.5, 1 M NaCl, 10% glycerol, 10 mM imidazole, and 1 mM DTT. Protein was eluted from resin with 300 mM imidazole in 20 mM HEPES pH 7.5, 400 mM NaCl, and 1 mM DTT. Fractions containing His-MBP-cGAS were pooled, concentrated, and dialyzed against 20 mM HEPES pH 7.5, 400 mM NaCl, 1 mM DTT, and then snap-frozen in aliquots for future use.

### Synthesis and purification of cGAMP

To enzymatically synthesize cGAMP, 1 μM purified cGAS was incubated with 50 mM Tris-HCl pH 7.4, 2 mM ATP, 2 mM GTP, 20 mM MgCl_2_, and 100 μg/mL herring testis DNA (Sigma) for 24 h. The reaction was then heated at 95 °C for 3 min and filtered through a 3-kDa filter. cGAMP was purified from the reaction mixture using a PLRP-S polymeric reversed phase preparatory column (100 Å, 8 μm, 300 × 25 mm; Agilent Technologies) on a preparatory HPLC (1260 Infinity LC system; Agilent Technologies) connected to UV-vis detector (ProStar; Agilent Technologies) and fraction collector (440-LC; Agilent Technologies). The flow rate was set to 25 mL/min. The mobile phase consisted of 10 mM triethylammonium acetate in water and acetonitrile. The mobile phase started as 2% acetonitrile for first 5 min. Acetonitrile was then ramped up to 30% from 5-20 min, then to 90% from 20-22 min, maintained at 90% from 22-25 min, and then ramped down to 2% from 25-28 min. Fractions containing cGAMP were lyophilized and resuspended in water. The concentration was determined by measuring absorbance at 280 nm.

### Cell culture

HEK 293T cells were maintained in DMEM (Cellgro) supplemented with 10% FBS (Atlanta Biologicals) and 1% penicillin-streptomycin (Gibco). All U937 cell lines were maintained in RPMI (Cellgro) supplemented with 10% heat-inactivated FBS (Atlanta Biologicals) and 1% penicillin-streptomycin (Gibco). CD14^+^ primary cells were maintained in RPMI (Cellgro) supplemented with 10% heat-inactivated FBS (Atlanta Biologicals), 1 mM sodium pyruvate (Gibco), 50 μM 2-mercaptoethanol (Sigma), and 10 ng/mL M-CSF (PeproTech). All cell lines were obtained from ATCC. All cells were maintained in a 5% CO_2_ incubator at 37 °C.

### U937 CRISPR knockout library generation

The methods used to create the U937 CRISPR knockout library line were previously described by Morgens et al. (2017). Briefly, a whole-genome library of exon-targeting sgRNAs were designed, with the goal of minimizing off-target effects and maximizing gene disruption. The top 10 sgRNA sequences for each gene were included in the library, along with thousands of safe-targeting and non-targeting negative controls. The library was cloned into a lentiviral vector, pMCB320, which also expresses mCherry and a puromycin resistance cassette. The cells were infected with the lentiviral library, and then selected with puromycin.

### CRISPR screen

The U937 CRISPR knockout library line was grown in 4 spinner flasks (1 L), with 2 flasks serving as untreated controls and 2 flasks receiving cGAMP treatment. Throughout the screen all of the samples were split daily to keep the cell density at 250 million cells per 500 mL, which corresponded to 1,000 cells per guide in the untreated samples. The experimental samples were treated daily with enough cGAMP to reduce cell viability by 50% as compared to the control samples. The initial treatment was 20 μM and was increased steadily over the 2-week screen, ultimately reaching 30 μM on the final treatment day. At the end of 2 weeks, the genomic DNA was extracted using a Qiagen Blood Maxi Kit. The library was sequenced using a NextSeq 500/550 Mid Output v2 kit (Illumina). The experimental and control samples were compared using casTLE (Morgens et al. 2016), available at https://bitbucket.org/dmorgens/castle. The algorithm determines the likely effect size for each gene, as well as the statistical significance of this effect.

### PBMC isolation

Buffy coat (Stanford Blood Center) was diluted 1:3 with PBS supplemented with 2 mM EDTA. Diluted buffy coat was layered on top of 50% Percoll (GE Healthcare) containing 140 mM NaCl and centrifuged at 600 x g for 30 min. The separated PBMC layer was collected and washed once with PBS and once with RPMI before MACS CD14^+^ isolation.

### CD14^+^ primary cell MACS Isolation

CD14^+^ cells in bulk PBMCs were labeled using CD14 MicroBeads (Miltenyi Biotec) and purified using a MACS LS Column on a MidiMACS Separator (Miltenyi Biotec) following the manufacturer’s instructions.

### *IFNβ* qPCR

For detection of *IFNβ* transcript, U937 cells (5 × 10^5^ cells in 1 mL) were first treated with 100 μM cGAMP for 6 h. Cells were then pelleted and lysed with 1 mL TRIzol (Invitrogen). Total RNA from cells was extracted following manufacturer’s instructions. This RNA was reverse transcribed in 20 μL reactions containing 500 ng total RNA, 100 pmol Random Hexamer Primers (Thermo Scientific), 0.5 mM dNTPs (NEB), 20 U RNaseOUT (Invitrogen), 1x Maxima RT Buffer (Thermo Scientific), and 200 U Maxima Reverse Transcriptase (Thermo Scientific). Reverse transcription reactions were incubated first for 10 min at 37 °C, then for 30 min at 50 °C. Reactions were then terminated by incubating for 5 min at 85 °C. To quantify transcript levels, 10 μL reactions were set up containing 1x GreenStar Master Mix (Bioneer), 10x ROX dye (Bioneer), 100 nM forward and reverse primers, and 0.7 μL of reverse transcription reactions. To determine Ct values, reactions were run on a ViiA 7 Real-Time PCR System (Applied Biosystems) using the following program: ramp up to 50 °C (1.6 °C/s) and incubate for 2 min, ramp up to 95 °C (1.6 °C/s) and incubate for 10 min; then 40 cycles of ramp up to 95 °C (1.6 °C/s) and incubate for 15 sec, ramp down to 60 °C (1.6 °C/s) and incubate for 1 min. The following sets of primers were used: *IFNβ* transcript 5’-AAACTCATGAGCAGTCTGCA-3’ (forward), 5’-AGGAGATCTTCAGTTTCGGAGG-3’ (reverse); ACTB transcript 5’-GGCATCCTCACCCTGAAGTA-3’ (forward), 5’-AGAGGCGTACAGGGATAGCA-3’ (reverse).

### IFN-β Protein Detection

U937 cells (5 × 10^5^ cells in 1 mL) were treated with 100 μM cGAMP for 6 h. Then, cells were pelleted and supernatant containing secreted IFN-β was collected. 20 μL of supernatant were added to 180 μL HEK-Blue IFN-α/β cells (InvivoGen) (2.8 × 10^5^ cells/mL) and incubated for 20 h in a 5% CO_2_ incubator at 37 °C. Supernatant from HEK-Blue IFN-α/β cells was then assayed for alkaline phosphatase activity using QUANTI-Blue (InvivoGen), following manufacturer’s instructions.

### Cell Viability Measurements

U937 cells were treated with the indicated concentrations of cGAMP for 16 hours. 100 μL of CellTiter-Glo (Promega) were added directly to 100 μL of cells. The cells were shaken gently for 2 min, and then allowed to rest at room temperature for 8 min. The luminescence was measured on a Tecan Spark with a 500 ms integration time.

### *SLC19A1*^-/-^ and *cGAS*^-/-^ cell line generation

LentiCRISPR v2 (Addgene) was used as the 3^rd^-generation lentiviral backbone for all knockout lines. The guide sequence targeting *SLC19A1* was 5’-GCACGAGAGAGAAGATGT-3’, and the guide sequence targeting *cGAS* was 5’-GGCTTCCGCACGGAATGCCA-3’. The guide sequences were cloned into the lentiviral backbone using the Lentiviral CRISPR Toolbox protocol from the Zhang Lab at MIT(Sanjana, Shalem, and Zhang 2014; Shalem et al. 2014). Lentiviral packaging plasmids (pHDM-G, pHDM-Hgmp2, pHDM-tat1b, and RC/CMV-rev1b) were purchased from Harvard Medical School. 500 ng of the lentiviral backbone plasmid containing the guide sequence and 500 ng of each of the packaging plasmids were transfected into HEK 293T cells using FuGENE 6 transfection reagent (Promega). The viral media was exchanged after 24 hours and was harvested after 48 hours. The virus-containing media was passed through a 0.45 μm filter and added to U937 cells in media containing 8 μg/mL polybrene (Sigma Aldrich). The cells were spun at 1000 x g for 1 hour, and then the cells were resuspended in fresh virus-free media. The cells were put under selection with 1 μg/mL puromycin (Sigma Aldrich) for 1 week.

### Electroporation of STING agonists

For some experiments, cells were pretreated with 500 μM methotrexate for 5 min. 1 × 10^6^ cells were pelleted and resuspended in 100 μL electroporation solution (90 mM Na_2_HPO_4_, 90 mM NaH_2_PO_4_, 5 mM KCl, 10 mM MgCl_2_, 10 mM sodium succinate) with the appropriate concentration of STING agonist. Cells were then transferred to a cuvette with a 0.2 cm electrode gap (Bio-Rad) and electroporated using program U-013 on a Nucleofector II device (Lonza). Following electroporation, cells were transferred to the appropriate culture media and cultured for 90 min - 2 h (depending on the experiment) before processing for Western blot analysis.

### SLC19A1 Overexpression

To prepare for transfection, 7 × 10^6^ HEK 293T cells were split into a 12-well plate the day before transfection. These cells were then transfected with 50 ng of pcDNA3-STING-HA and 500 ng of either pcDNA3-FLAG-HA or pcDNA3-FLAG-HA-SLC19A1 using FuGENE 6 transfection reagent (Promega). After 24 h, transfected cells were treated with 100 μM cGAMP for 2 h. Following this, cells were directly lysed in Laemmli Sample Buffer and run on an SDS-PAGE gel for Western blot analysis.

### [^3^H]-Methotrexate uptake assay

2-4 × 10^6^ cells were treated with 17.5 nM [^3^H]-methotrexate in MHS buffer (20 mM HEPES, 225 mM sucrose, pH to 7.4 with MgO) for 5 min. Cold PBS was then added to cells to halt import. Following two more washes in cold PBS, cells were lysed in 500 μL 200 mM NaOH and heated at 65 °C for 45 min to completely dissolve the lysate. Then, 400 μL of cell lysate was used to count ^3^H signal on a scintillation counter and 25 μL were used in a bicinchoninic acid assay (Thermo Scientific) to quantify protein levels for normalization.

### Statistical analysis

All statistical analyses were performed using GraphPad Prism 7.4. For [^3^H]-methotrexate import in U937 cell lines, p values were calculated using an unpaired t-test assuming a Gaussian distribution. For all experiments involving Western blots, densitometric measurements of protein bands were made using ImageJ 1.51g. Measurements of total IRF3 or tubulin bands were used to normalize samples that were run on the same gel, and p values were calculated using a ratio paired t-test assuming a Gaussian distribution.

## Acknowledgements

We thank J. Carozza, K. Shaw, and S. L. Ergun for synthesizing cGAMP. We also thank all Li Lab members for their insightful comments and discussion throughout the course of this study. A.C. was supported by NIH 2T32GM007365-42. C.R. was supported by NIH 5T32GM007276. This work is supported by the National Institute of Health (5R00CA19089 and DP2CA228044 to L.L., and DP2HD084069 to M.B.).

## Competing Interests

The authors declare no competing interests.

