## Supplemental Figures for "SLC19A1 is an importer of the immunotransmitter cGAMP"

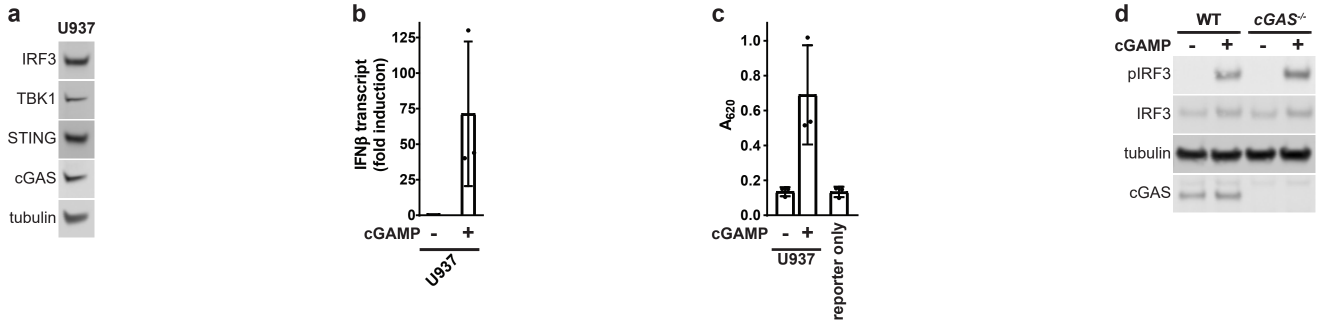


e

| | LD$_{50}$ Screen | | | LD$_{30}$ Screen | | |
|---|---|---|---|---|---|---|
| | casTLE Effect | casTLE Score | casTLE p-value (Maximum Estimate) | casTLE Effect | casTLE Score | casTLE p-value (Maximum Estimate) |
| **TMEM173** | 8.6 | 537 | $9.08 \times 10^{-6}$ | 4.8 | 459 | $1.00 \times 10^{-5}$ |
| **IRF3** | 8 | 507 | $9.08 \times 10^{-6}$ | 4.7 | 372 | $1.00 \times 10^{-5}$ |
| **TBK1** | 7.6 | 465 | $9.08 \times 10^{-6}$ | 4.3 | 308 | $1.00 \times 10^{-5}$ |
| **SLC19A1** | 4.8 | 187 | $9.08 \times 10^{-6}$ | 3.5 | 226 | $1.00 \times 10^{-5}$ |

**Supplementary Figure 1 - A genetic screen identifies putative components of the extracellular cGAMP-STING pathway. (a)** TBK1, IRF3, STING, and cGAS levels in U937 cells were probed using Western blots. All uncropped blots are available in the Extended Data. **(b)** Production of IFN-β mRNA in response to cGAMP treatment. U937 cells were treated with 100 μM cGAMP for 6 h. Total RNA was isolated and fold induction of IFN-β over untreated cells was quantified by RT-qPCR. Data shown as mean ± SD (n = 3 biological replicates). **(c)** Production of IFN-β protein in response to cGAMP treatment. U937 cells were treated with 100 μM cGAMP for 6 h. Supernatant containing secreted IFN-β was then collected and added to HEK-Blue IFN-α/β reporter cells to quantify IFN-β protein levels. Data shown as mean ± SD (n = 3 biological replicates). **(d)** Effect of cGAS on extracellular cGAMP signaling. U937 WT and *cGAS*$^{-/-}$ cells were treated with 100 μM cGAMP for 90 min and probed for pIRF3. **(e)** Table showing the casTLE effects, scores, and p-values for the STING pathway genes *TMEM173*, *IRF3*, and *TBK1*, as well as the candidate cGAMP importer *SLC19A1*. All values were calculated using the methods described in Morgens *et al* (2016).

**a**

| clone | allele | mutations in *SLC19A1* gene |
| --- | --- | --- |
| *SLC19A1*-/- #1 | 1 | Δ10548-10554 |
| | 2 | Δ10548-10554 |
| *SLC19A1*-/- #2 | 1 | A10292G; Δ10548-10557 |
| | 2 | G10226C; Δ10227-10556 |

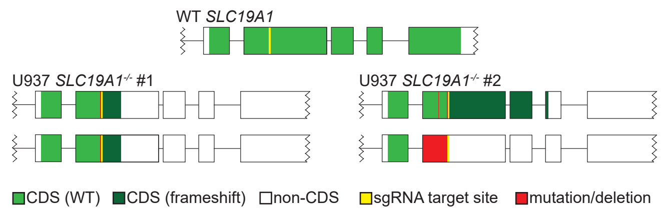


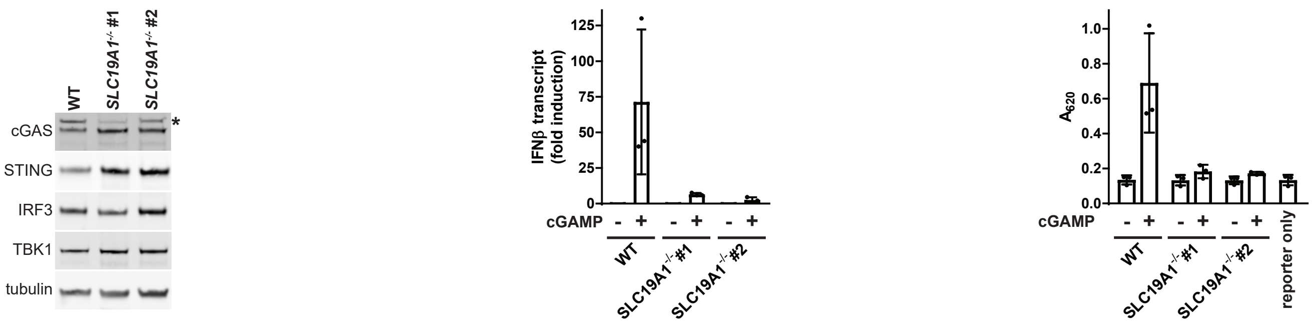


**Supplementary Figure 2 - SLC19A1 is essential for robust extracellular cGAMP signaling in U937 cells. (a)** Mutations at SLC19A1 loci in U937 *SLC19A1*-/- cell lines. Genomic DNA was isolated from U937 *SLC19A1*-/- cells and 800 bp of DNA flanking the *SLC19A1* sgRNA target site was sequenced. Note that both alleles in U937 *SLC19A1*-/- #1 contain the same mutation, whereas the alleles in U937 *SLC19A1*-/- #2 contain different mutations. **(b)** Diagrams depicting the predicted coding DNA sequence of *SLC19A1* alleles in *SLC19A1*-/- cells. Boxes represent coding exons of *SLC19A1* and horizontal lines represent introns. **(c)** TBK1, IRF3, STING, and cGAS levels in U937 WT and *SLC19A1*-/- cells were assayed by Western blot. * indicates nonspecific band. **(d)** Role of SLC19A1 in IFN-β mRNA production in response to cGAMP treatment. U937 WT and *SLC19A1*-/- cells were treated with 100 μM cGAMP for 6 h. Total RNA was isolated and fold induction of IFN-β over untreated cells was quantified by RT-qPCR. Data shown as mean ± SD (n = 3 biological replicates). **(e)** Role of SLC19A1 in IFN-β protein production in response to cGAMP treatment. U937 WT and *SLC19A1*-/- cells were treated with 100 μM cGAMP for 6 h. Supernatant containing secreted IFN-β was then collected and added to HEK-Blue IFN-α/β reporter cells to quantify IFN-β protein levels. Data shown as mean ± SD (n = 3 biological replicates).

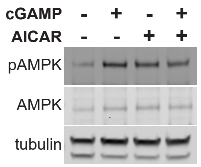


**Supplementary Figure 4 -SLC19A1 is a direct cGAMP importer.** Combining AICAR and cGAMP treatment does not cause increased pAMPK. U937 cells were pretreated with 1 mM AICAR for 15 min., and then treated with cGAMP for 90 min.

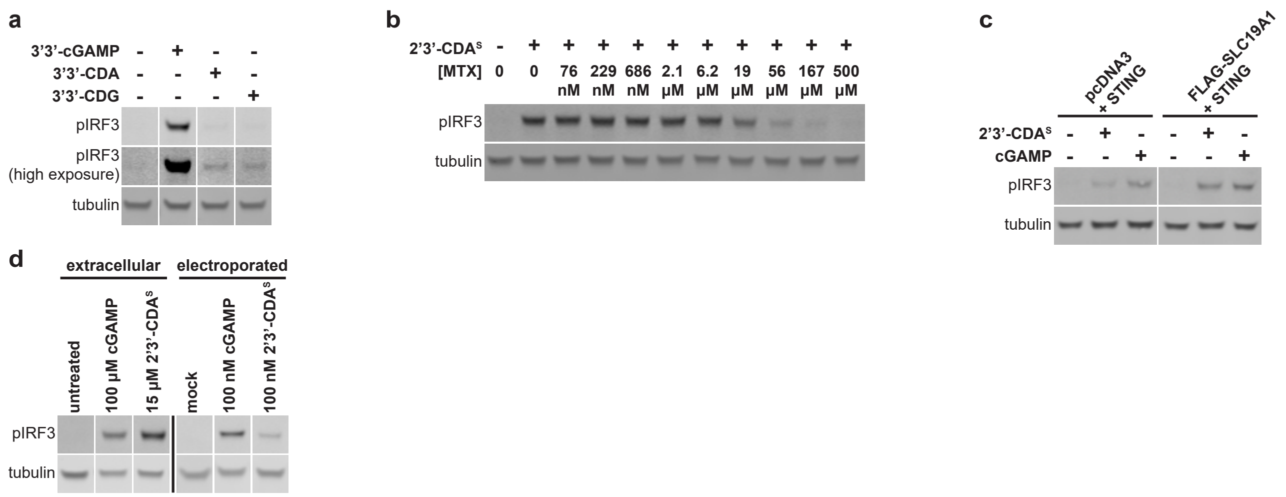


**Supplementary Figure 5 -SLC19A1 imports bacterial and synthetic cyclic dinucleotides, including 2'3'-CDA$^S$. (a)** Extracellular 3'3'-CDN signaling in U937 cells. U937 WT cells were treated with either 200 μM 3'3'-cGAMP, 400 μM 3'3'-CDA, or 400 μM 3'3'-CDG for 3 h. **(b)** Dose dependent effect of MTX on extracellular 2'3'-CDA$^S$ signaling. U937 cells were treated with 15 μM 2'3'-CDA$^S$ in the presence of various concentrations of MTX for 2 h. **(c)** Effect of SLC19A1 overexpression on extracellular 2'3'-CDA$^S$ signaling. HEK 293T cells were transfected with pcDNA3-STING-HA and either an empty pcDNA3-FLAG-HA vector or pcDNA3-FLAG-HA-SLC19A1. Transfected cells were treated 24 h after infection with 15 μM 2'3'-CDA$^S$ for 2 h. **(d)** Comparison of extracellular and intracellular response to cGAMP and 2'3'-CDA$^S$. U937 cells were treated with either 100 μM extracellular cGAMP or 15 μM 2'3'-CDA$^S$, or electroporated with 100 nM cGAMP or 2'3'-CDA$^S$ for 2 h.
