## Extended Figures for "SLC19A1 is an importer of the immunotransmitter cGAMP"

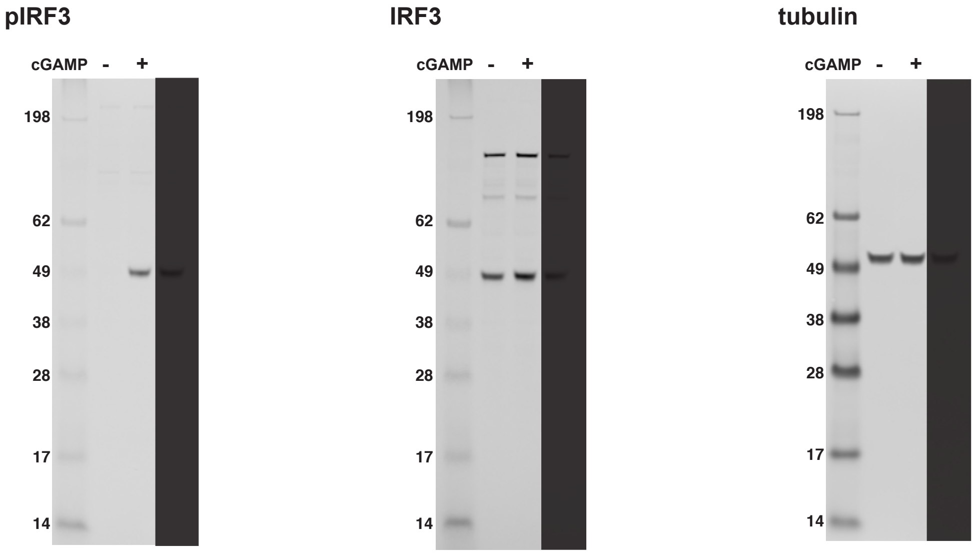


**Extended Figure 1a -** Uncropped Western blots for Figure 1a. Lanes not relevant to experiment are shaded.

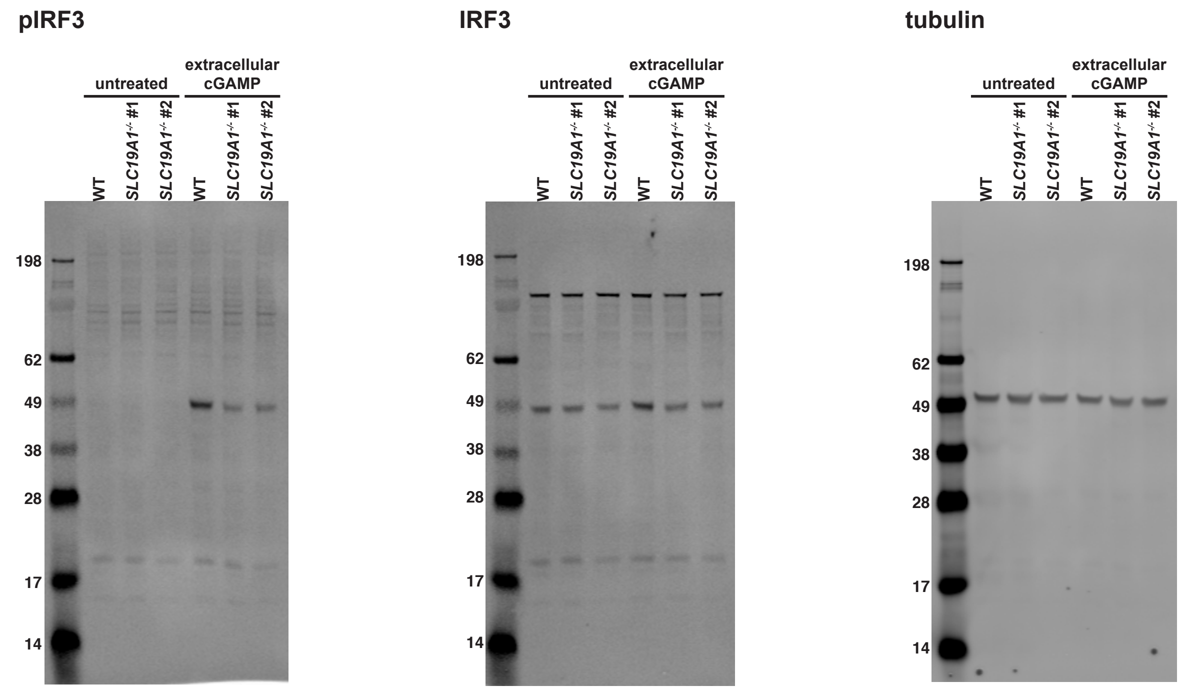


**Extended Figure 2c -** Uncropped Western blots for Figure 2c.

**pIRF3**

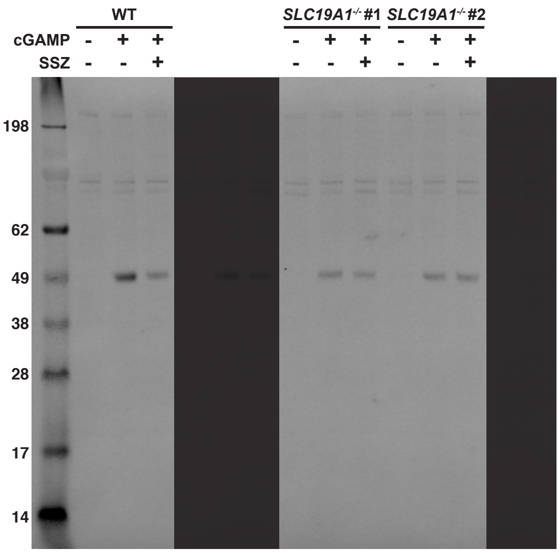


**IRF3**

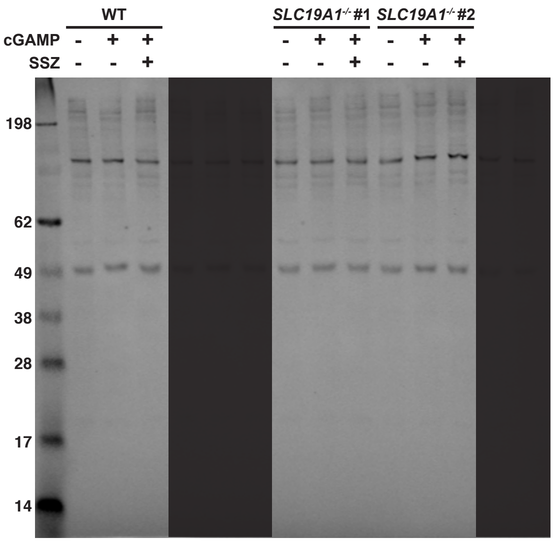


**tubulin**

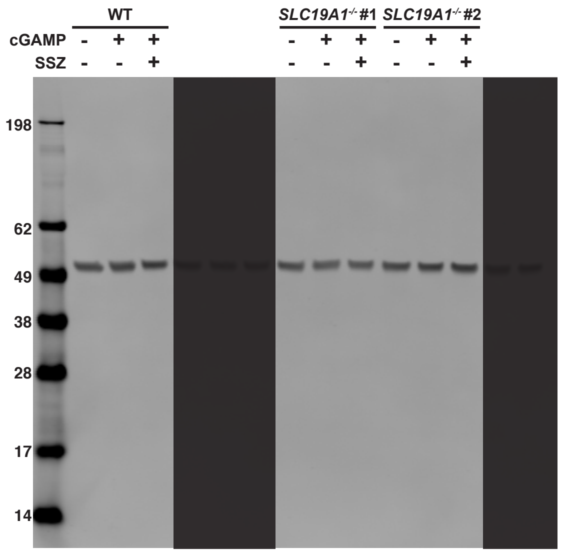


**Extended Figure 2d -** Uncropped Western blots for Figure 2d. Lanes not relevant to experiment are shaded.

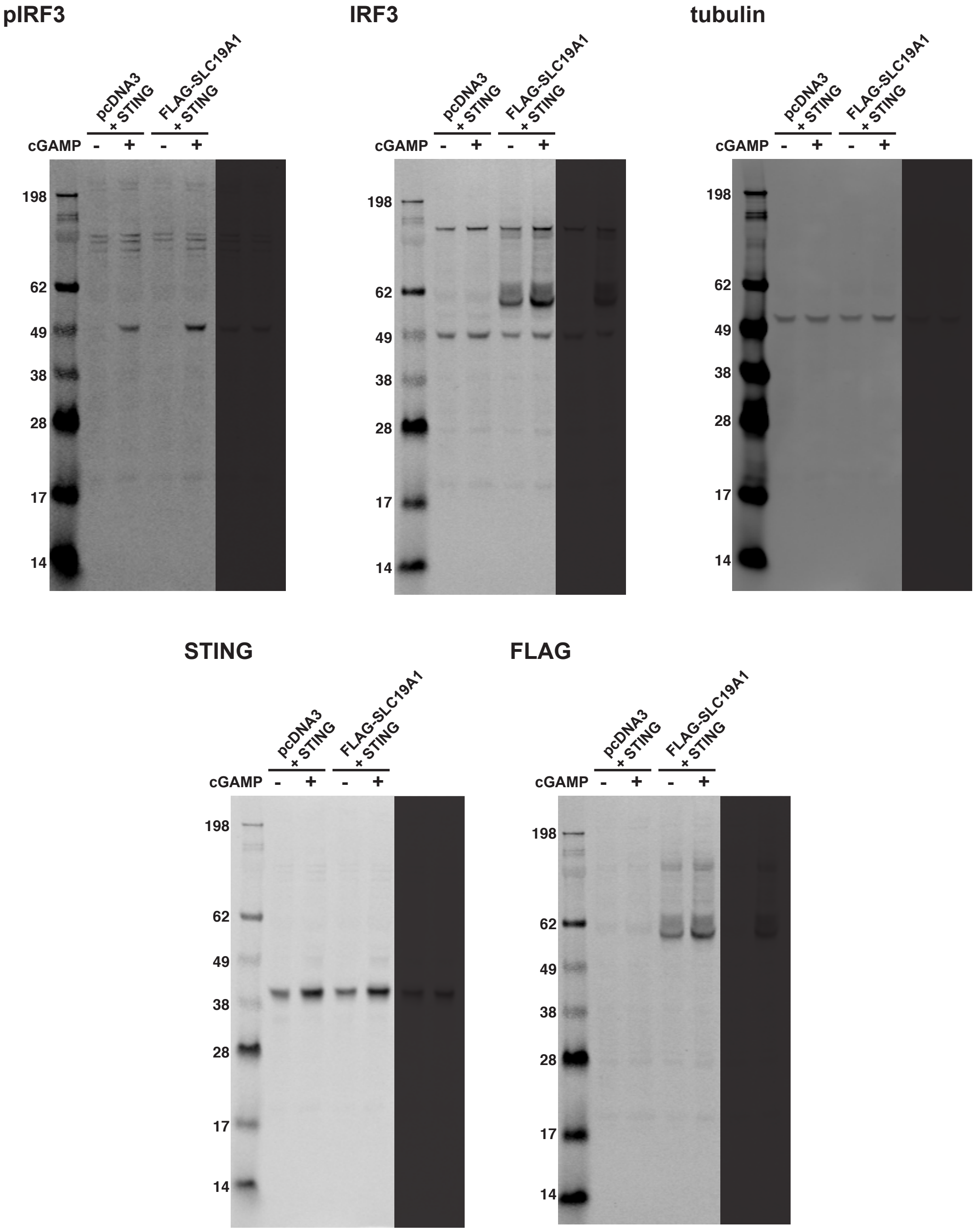


**Extended Figure 2e -** Uncropped Western blots for Figure 2e. Lanes not relevant to experiment are shaded.

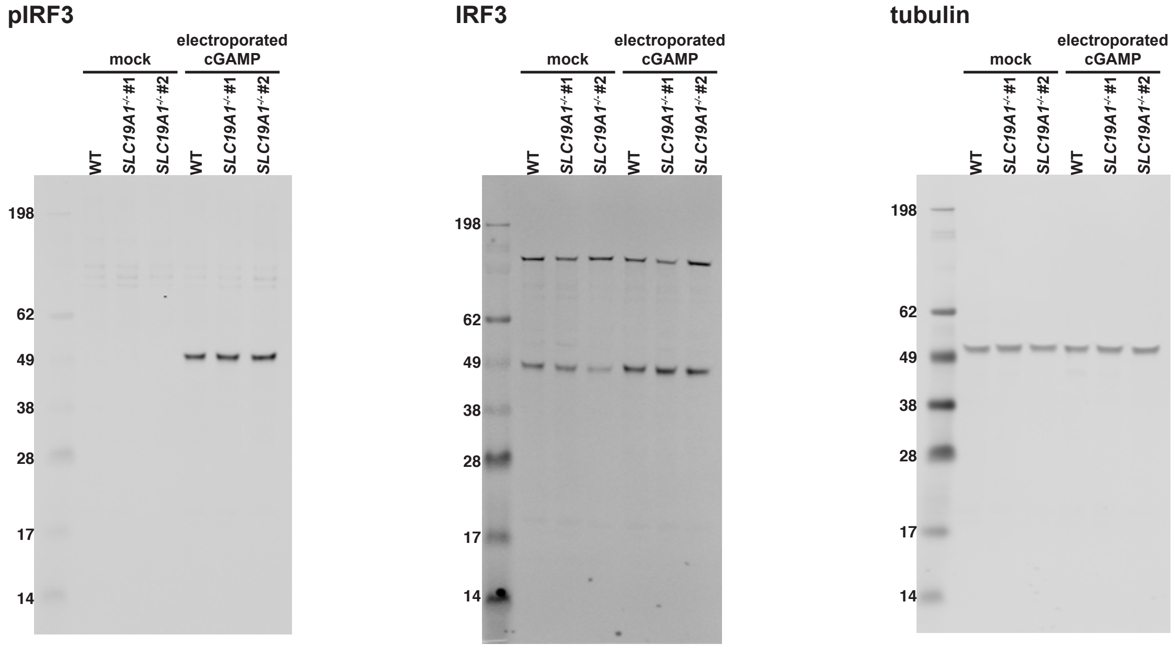


**Extended Figure 3b -** Uncropped Western blots for Figure 3b.

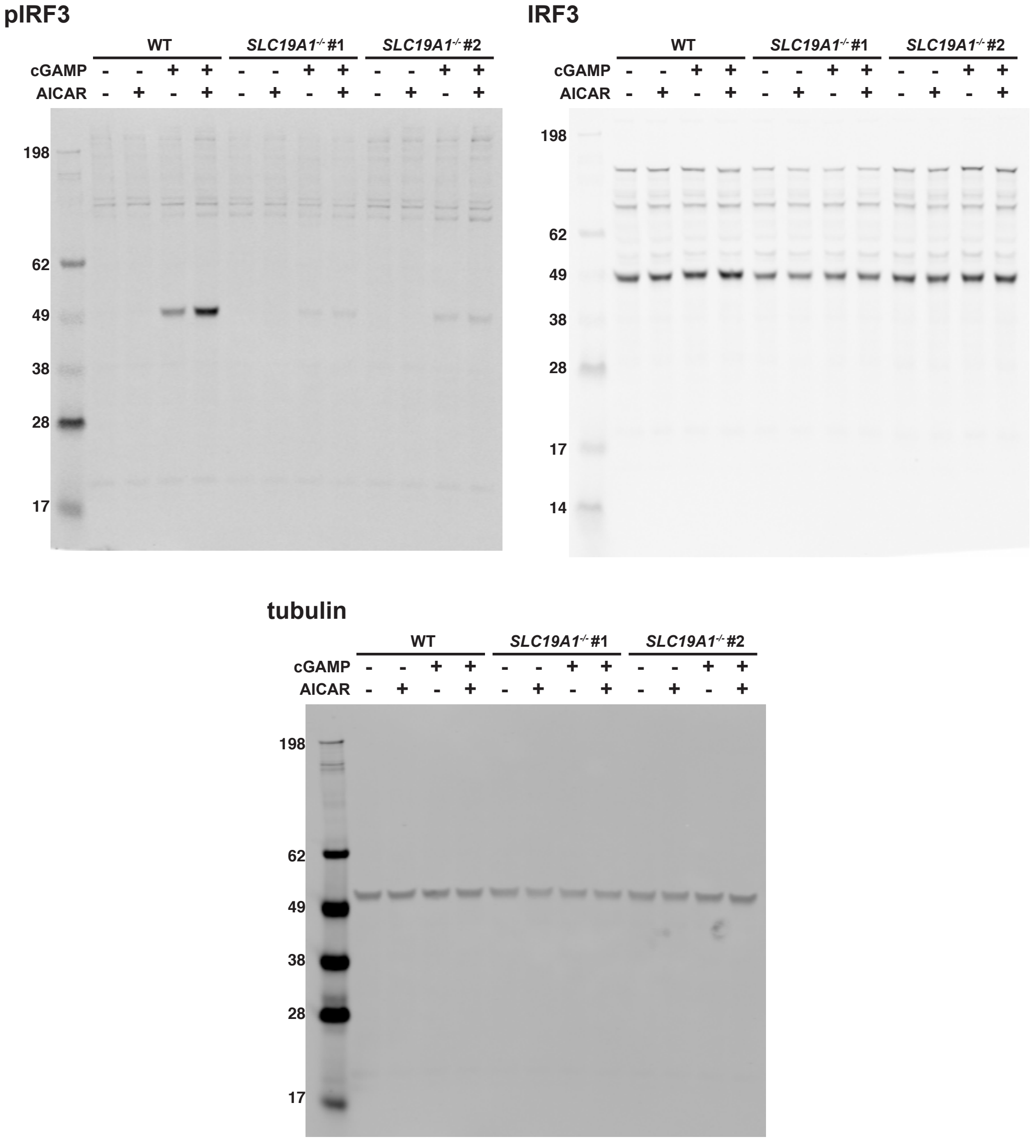


**Extended Figure 4a -** Uncropped Western blots for Figure 4a.

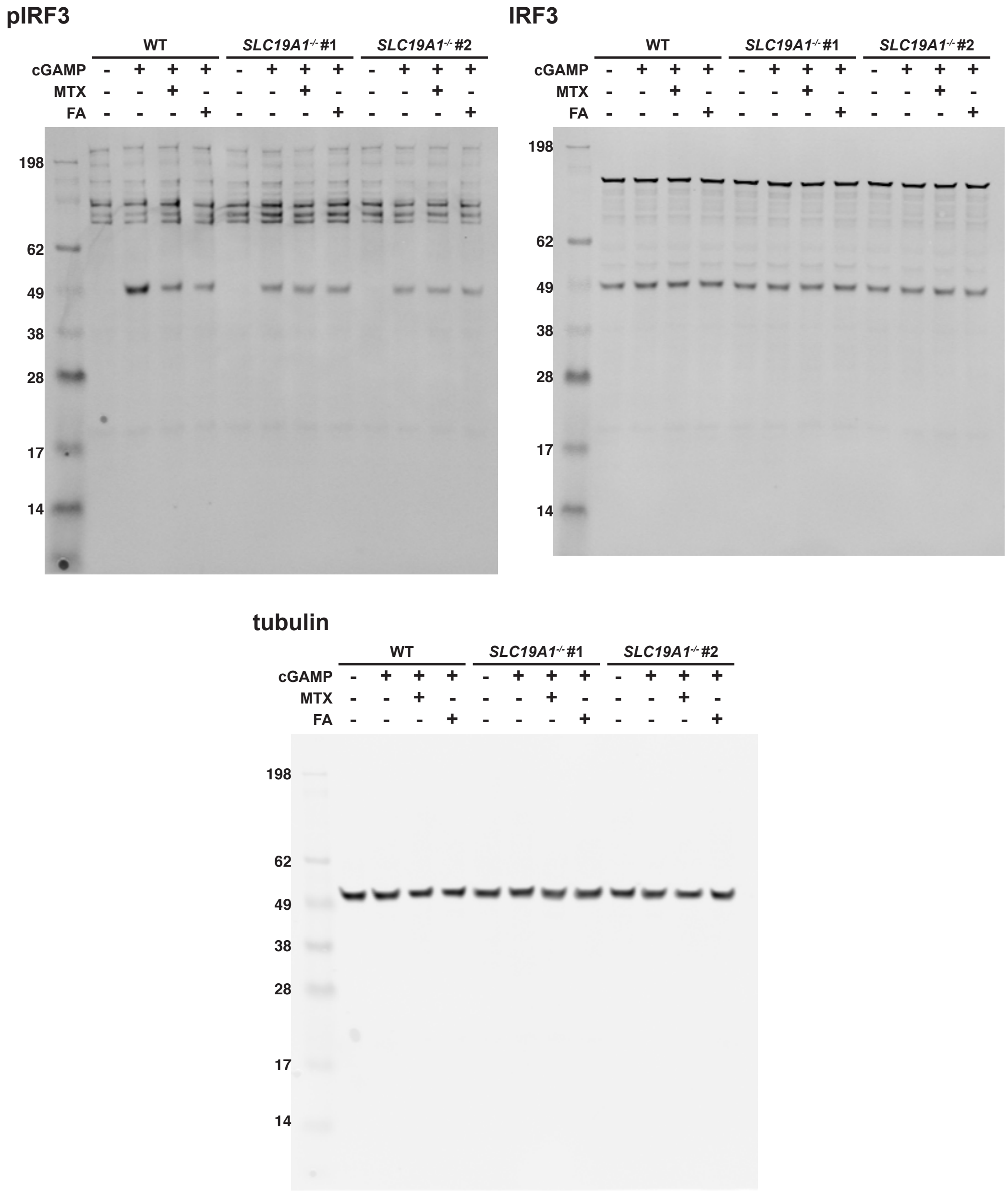


**Extended Figure 4b -** Uncropped Western blots for Figure 4b.

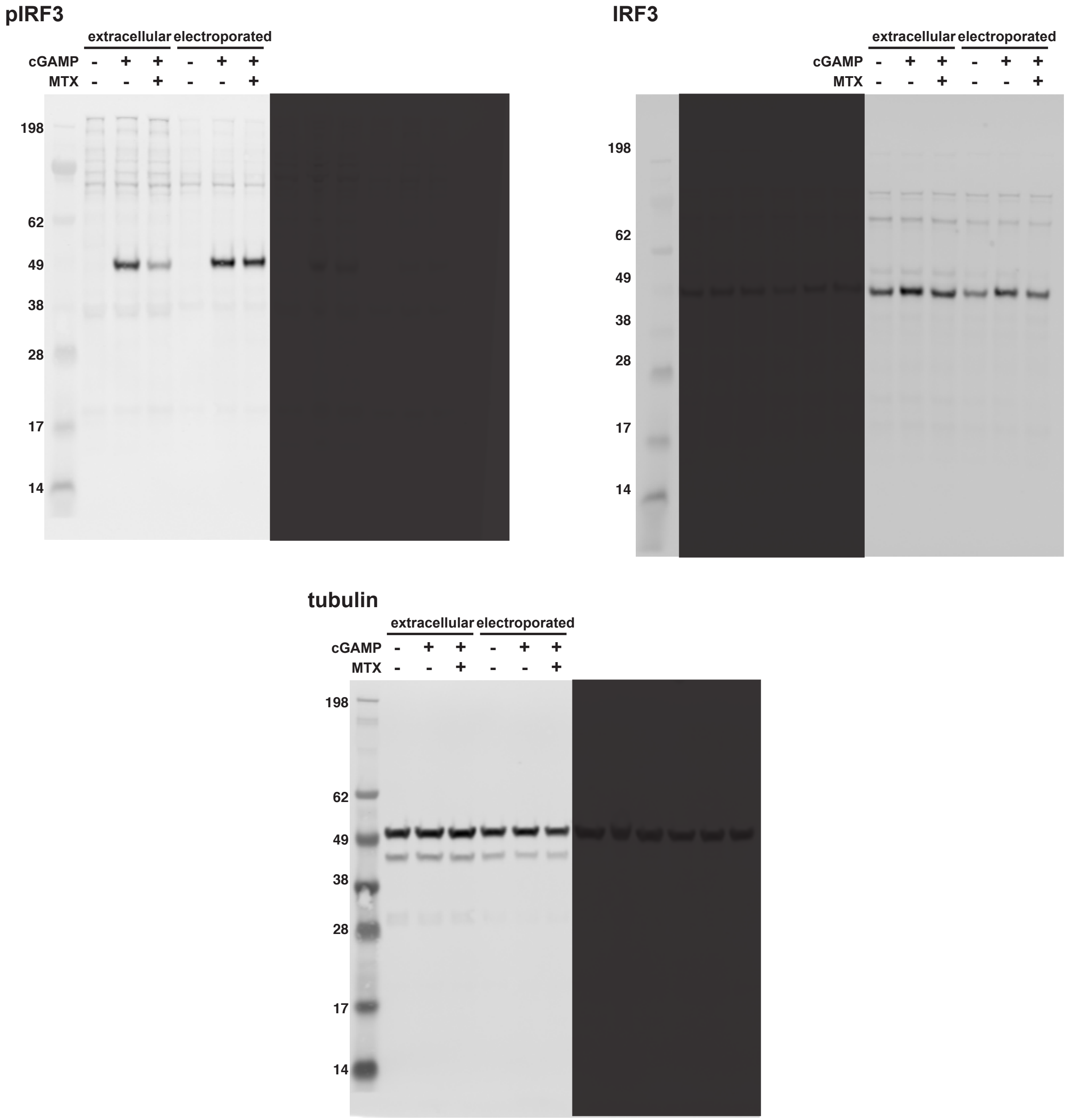


**Extended Figure 4c -** Uncropped Western blots for Figure 4c. Lanes not relevant to experiment are shaded.

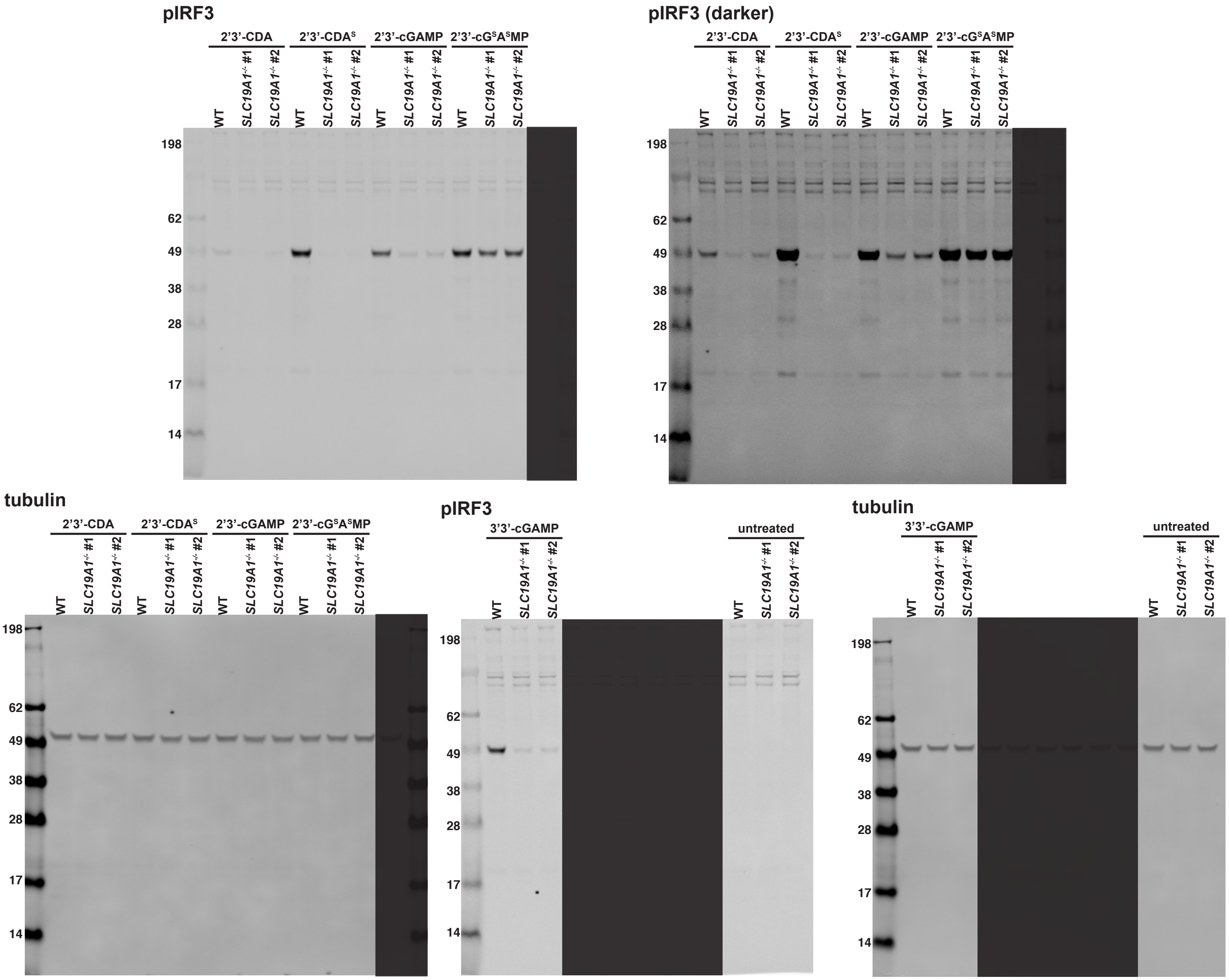


**Extended Figure 5b -** Uncropped Western blots for Figure 5b. Lanes not relevant to experiment are shaded.

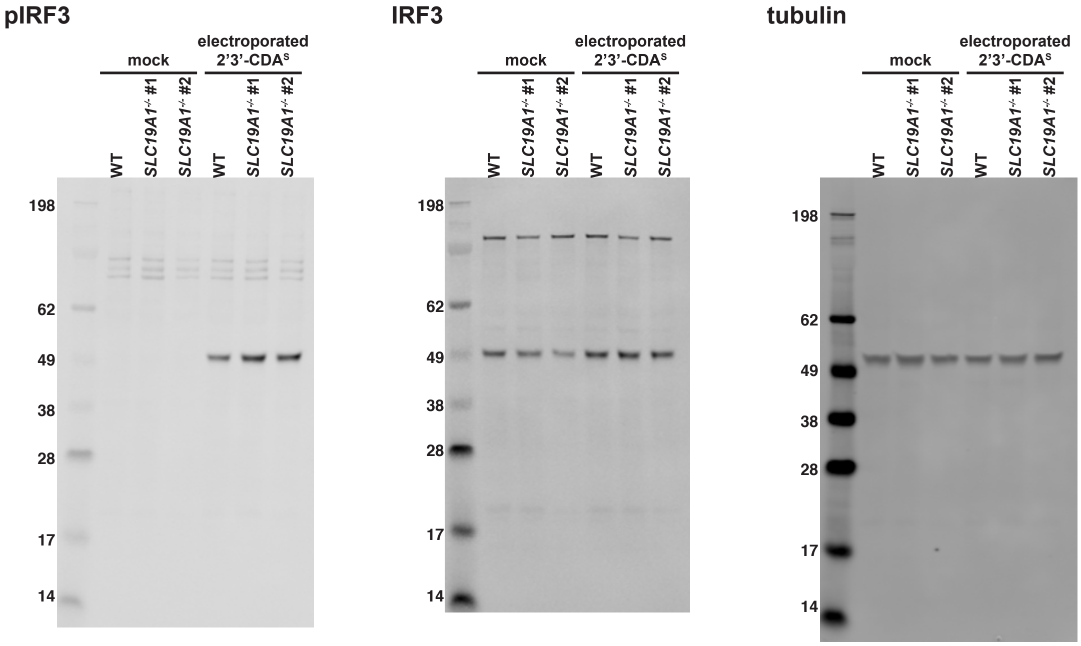


**Extended Figure 5c -** Uncropped Western blots for Figure 5c.

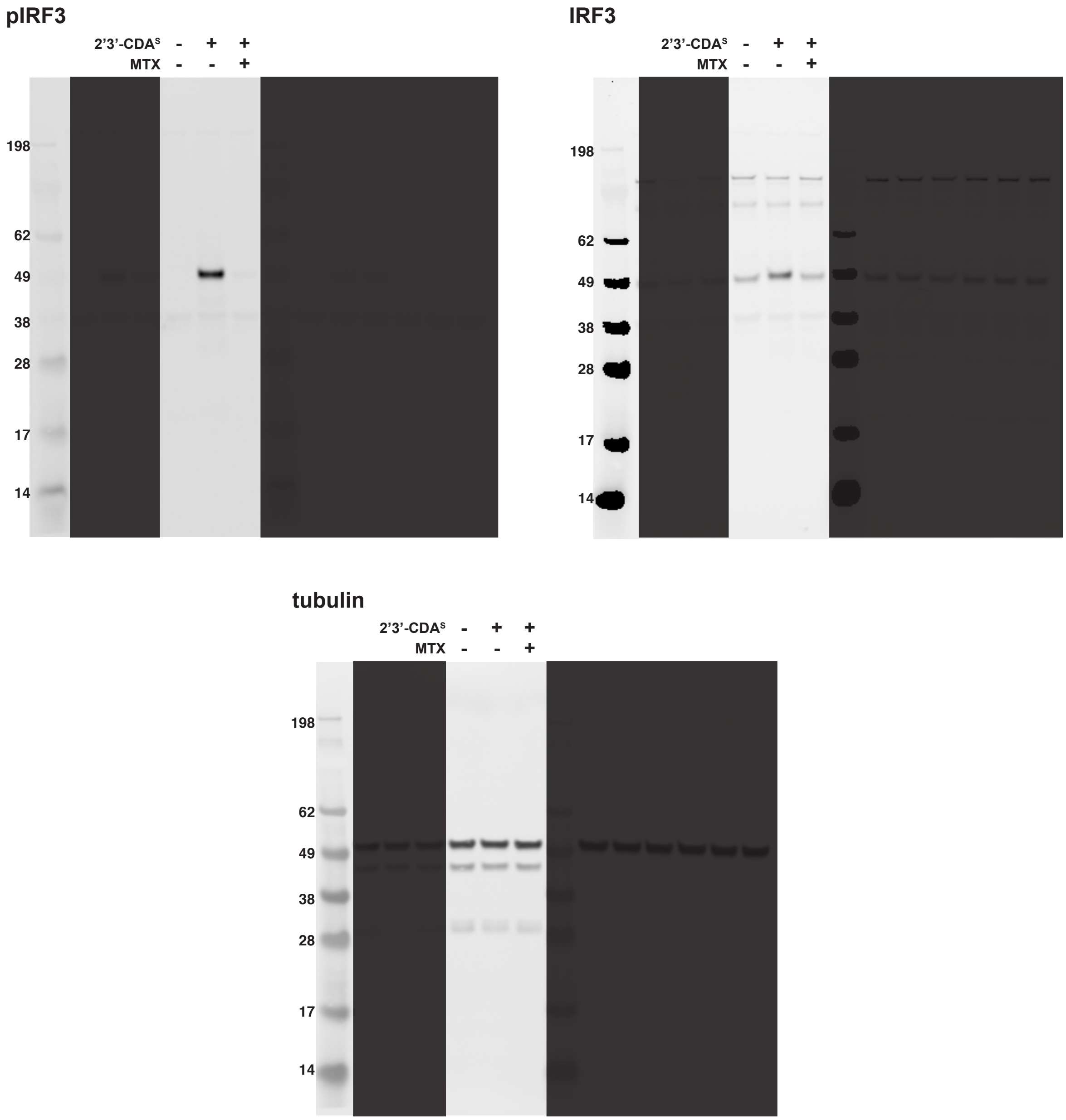


**Extended Figure 5d -** Uncropped Western blots for Figure 5d. Lanes not relevant to experiment are shaded.

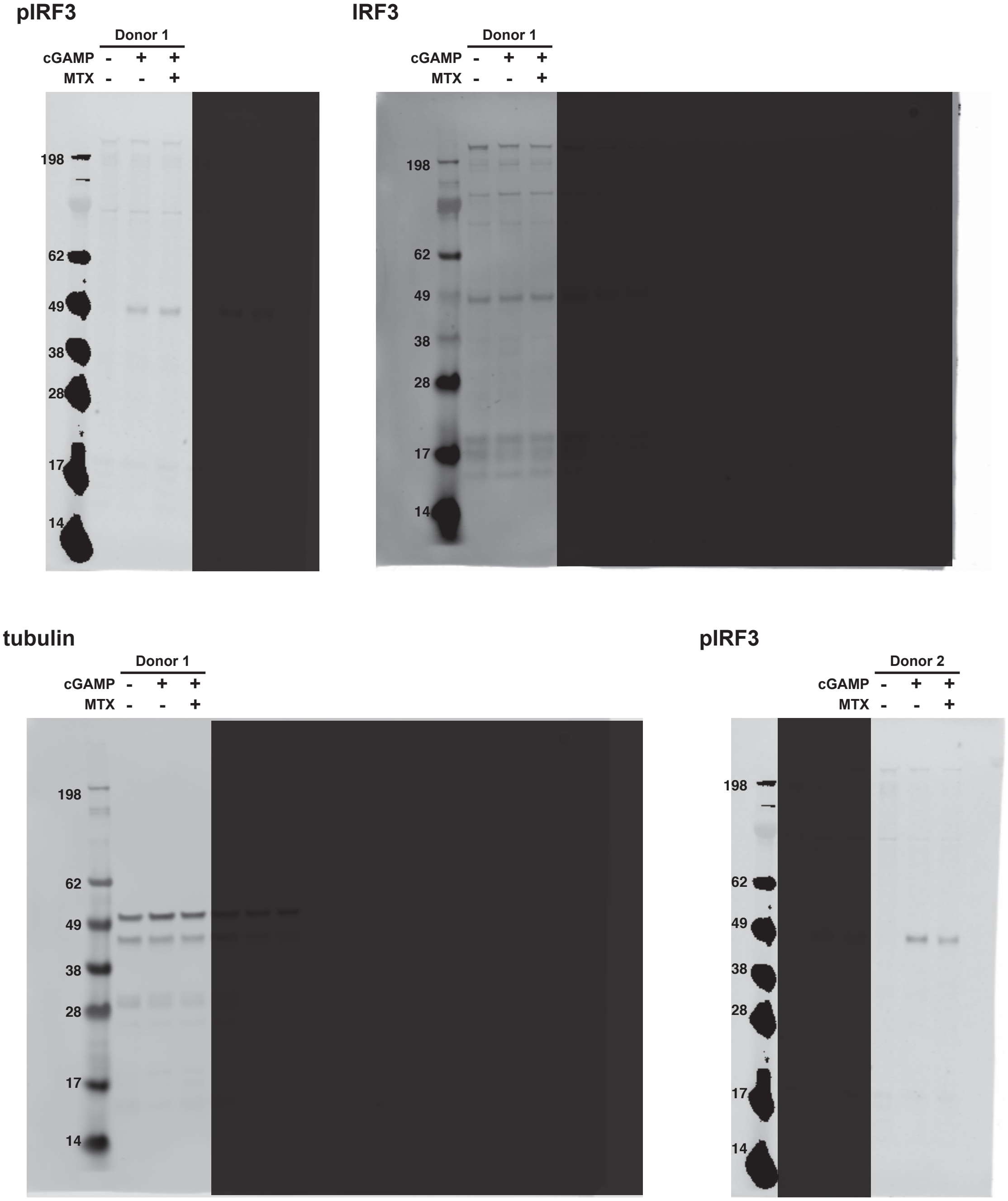


**Extended Figure 5e -** Uncropped Western blots for Figure 5e. Lanes not relevant to experiment are shaded.

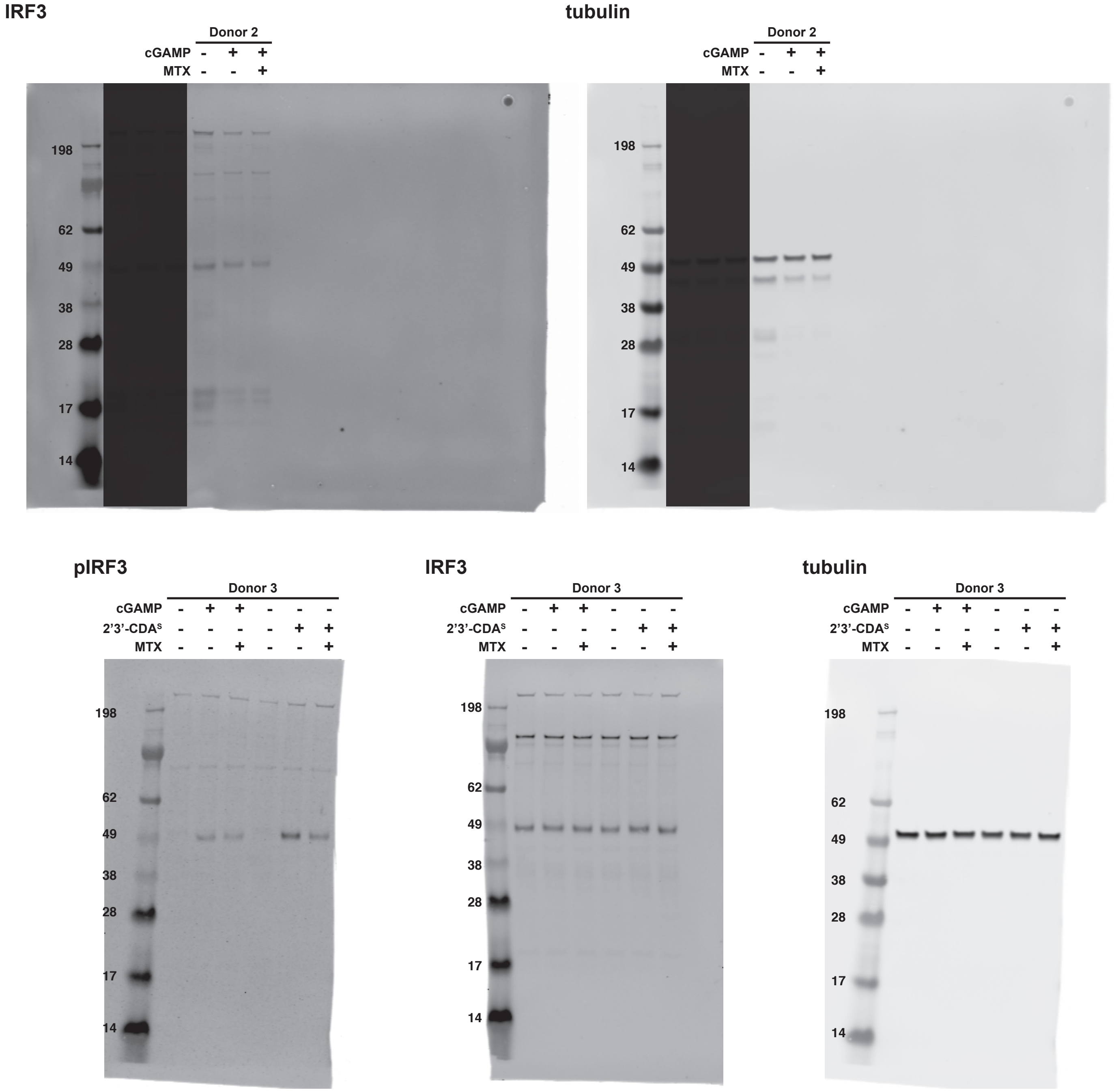


**Extended Figure 5e (cont.) -** Uncropped Western blots for Figure 5e. Lanes not relevant to experiment are shaded.

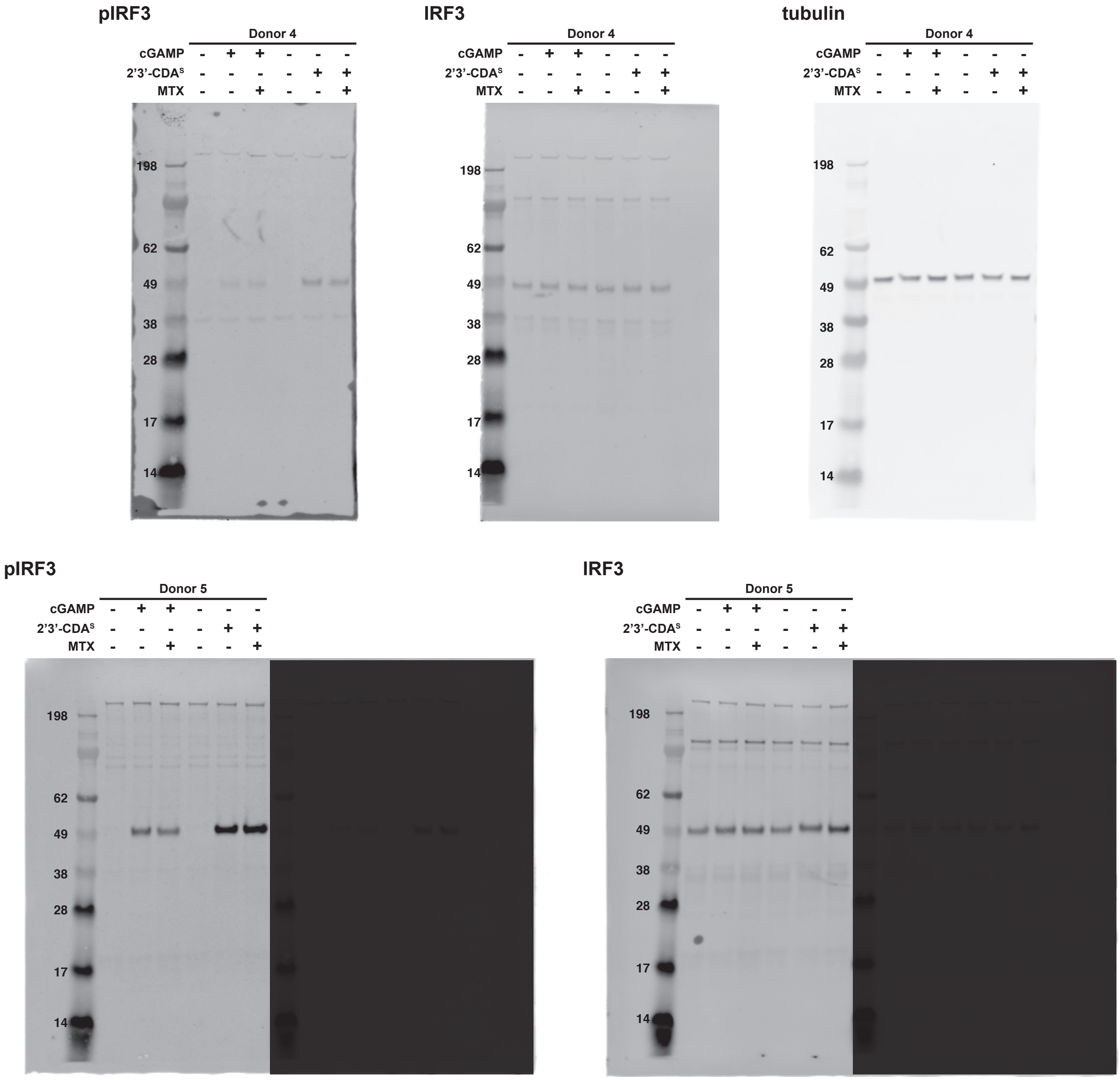


**Extended Figure 5e (cont.) -** Uncropped Western blots for Figure 5e. Lanes not relevant to experiment are shaded.

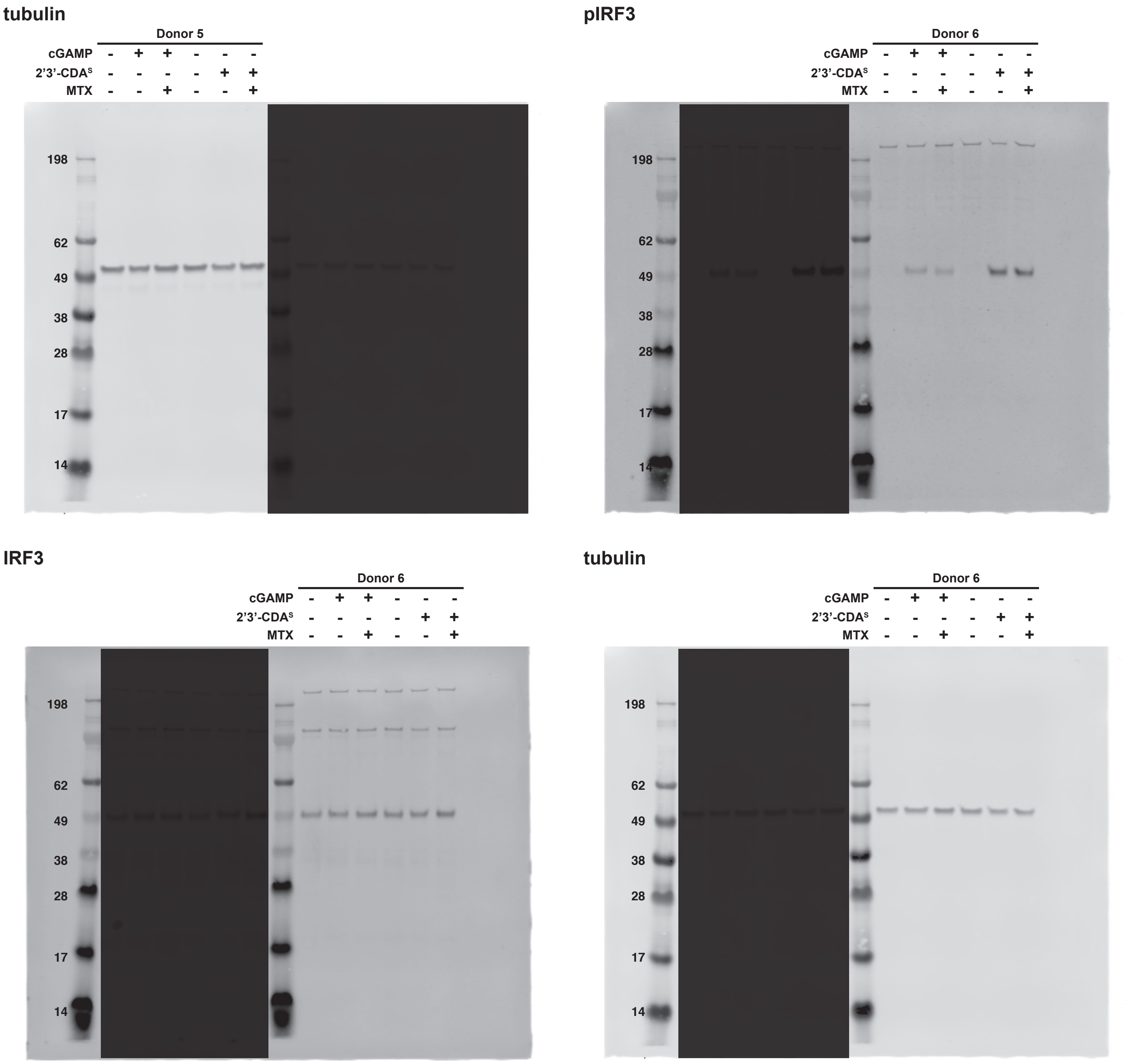


**Extended Figure 5e (cont.) -** Uncropped Western blots for Figure 5e. Lanes not relevant to experiment are shaded.

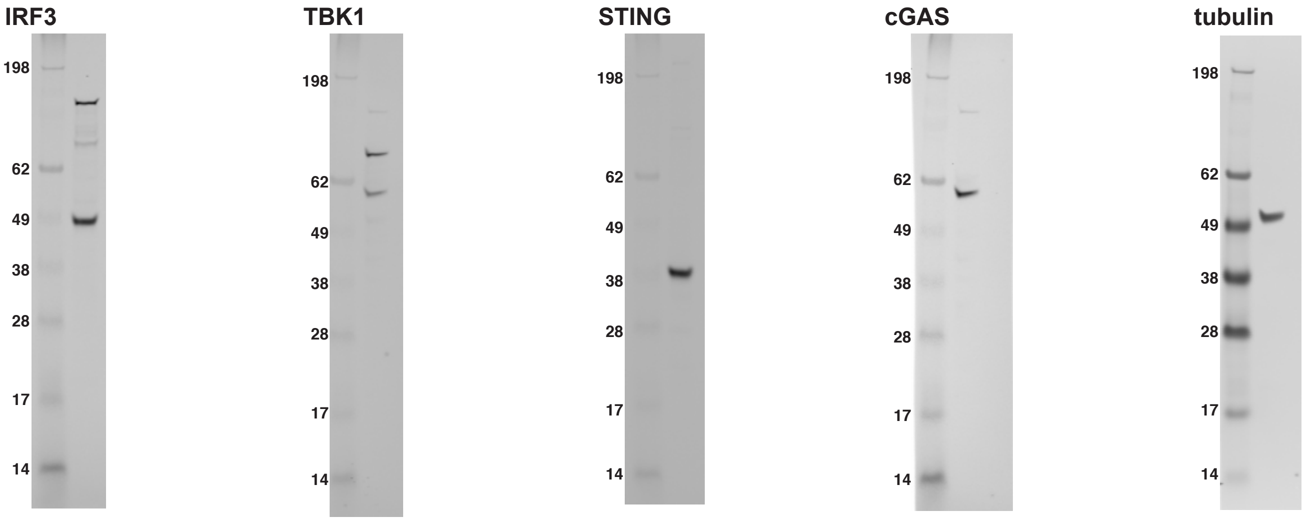


**Extended Supplementary Figure 1a -** Uncropped Western blots for Supplementary Figure 1a.

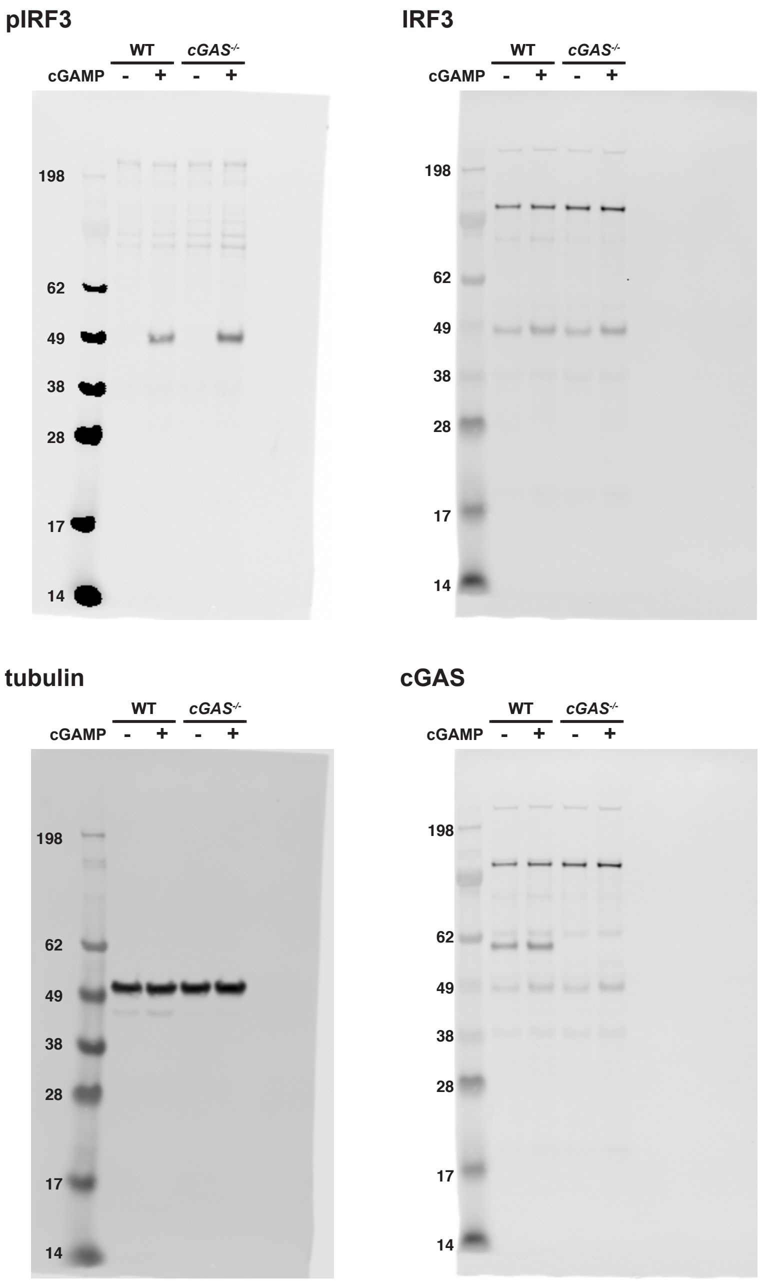


**Extended Supplementary Figure 1d -** Uncropped Western blots for Supplementary Figure 1d.

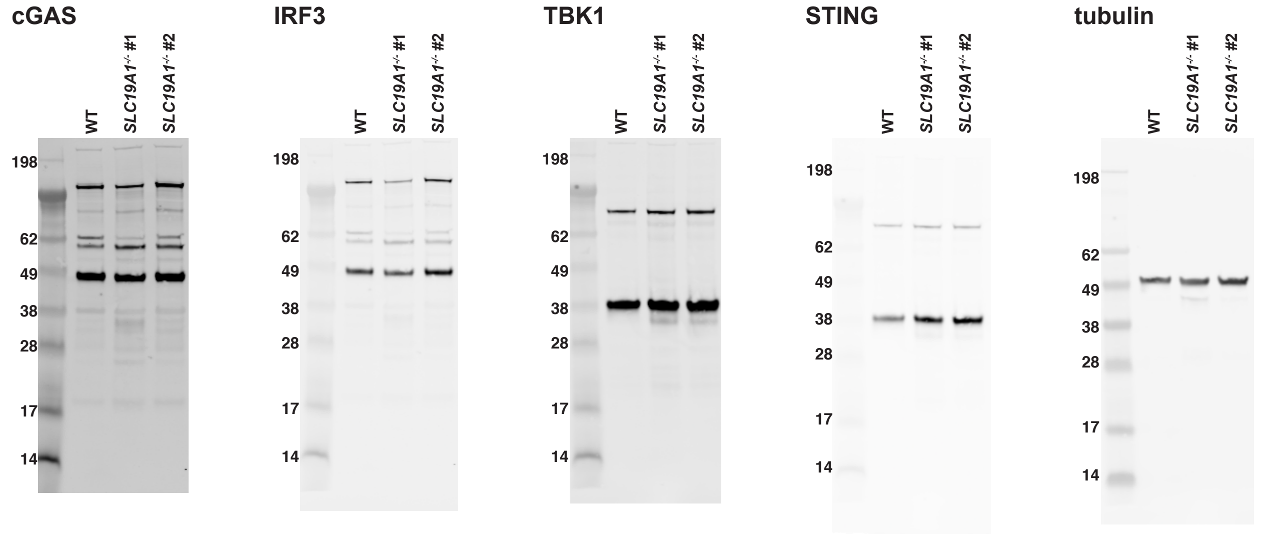


**Extended Supplementary Figure 2c -** Uncropped Western blots for Supplementary Figure 2c.

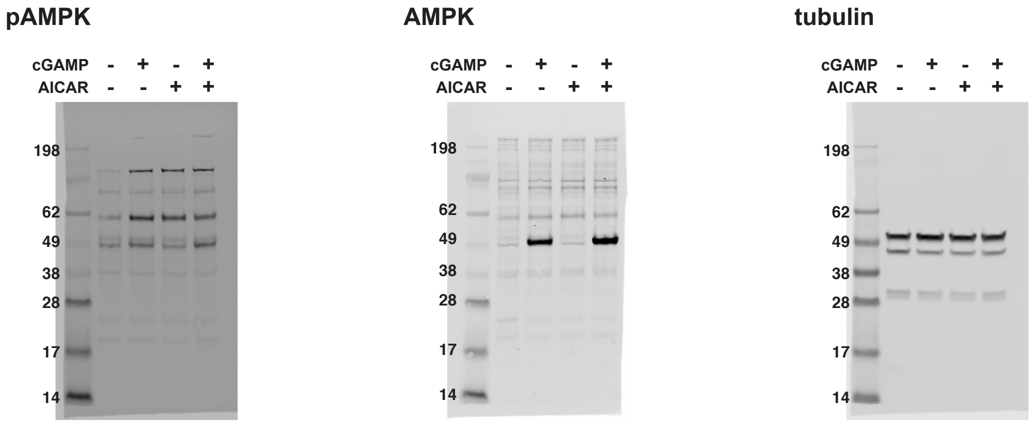


**Extended Supplementary Figure 4 -** Uncropped Western blots for Supplementary Figure 4.

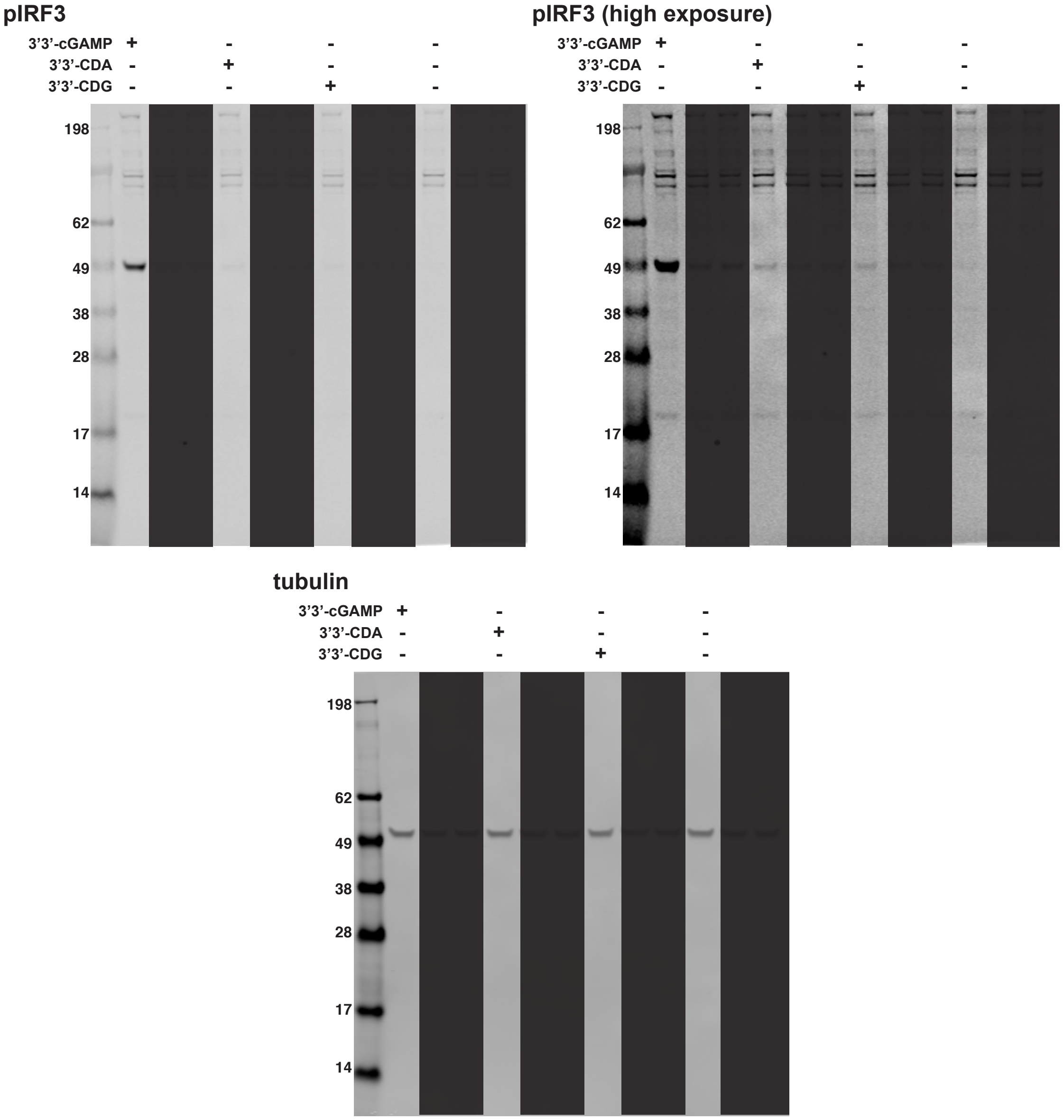


**Extended Supplementary Figure 5a -** Uncropped Western blots for Supplementary Figure 5a. Lanes not relevant to experiment are shaded.

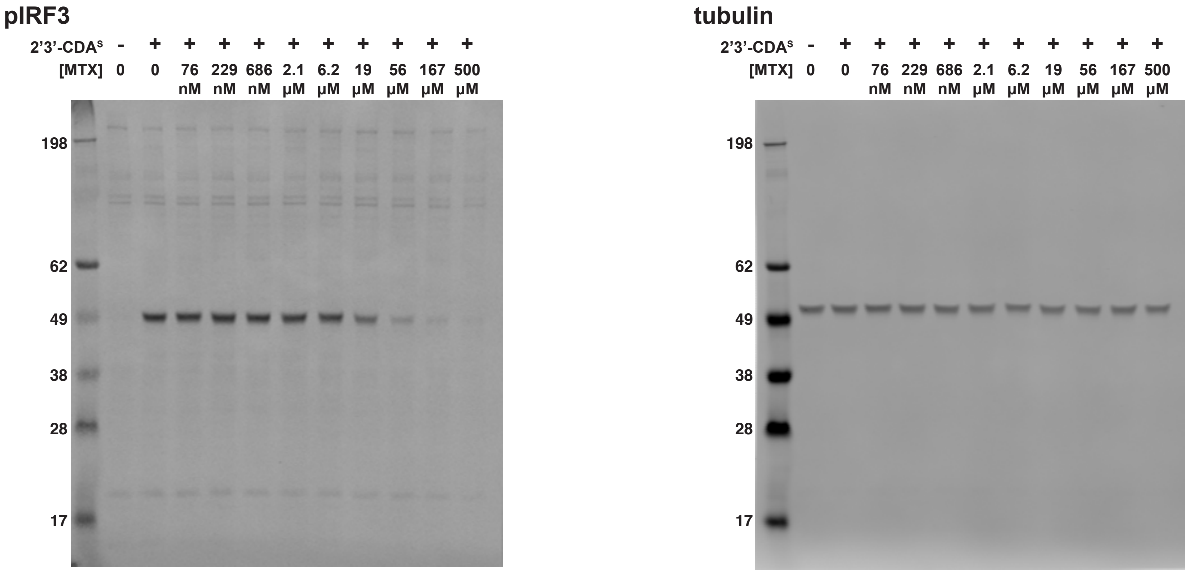


**Extended Supplementary Figure 5b -** Uncropped Western blots for Supplementary Figure 5b.

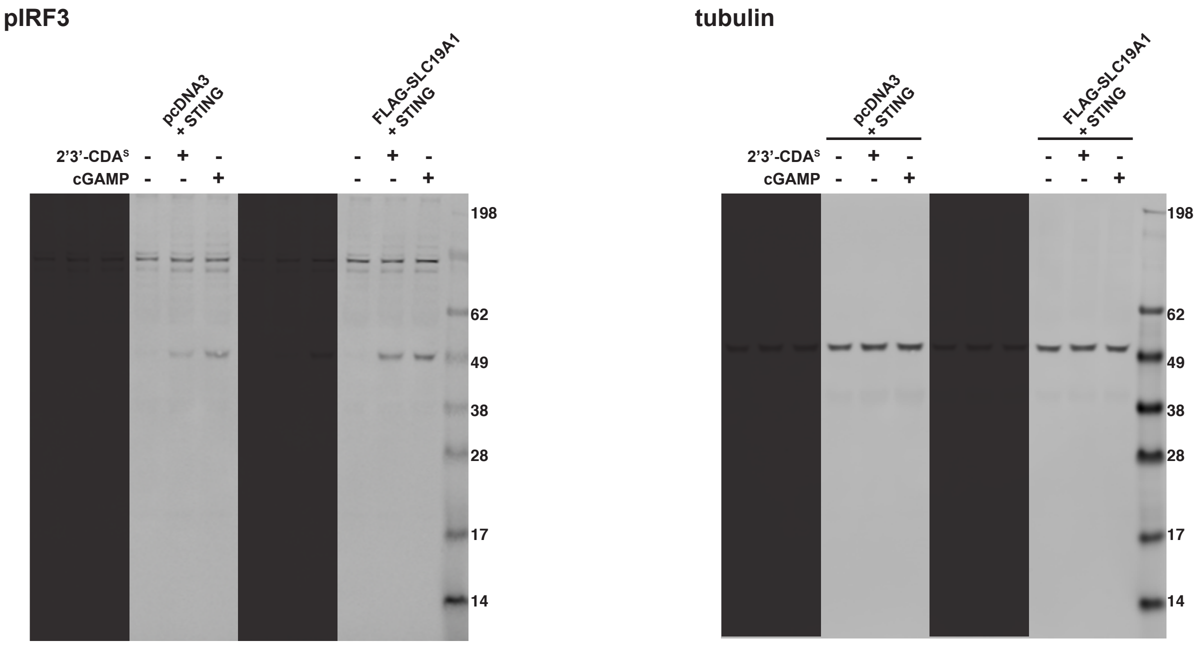


**Extended Supplementary Figure 5c -** Uncropped Western blots for Supplementary Figure 5c. Lanes not relevant to experiment are shaded.

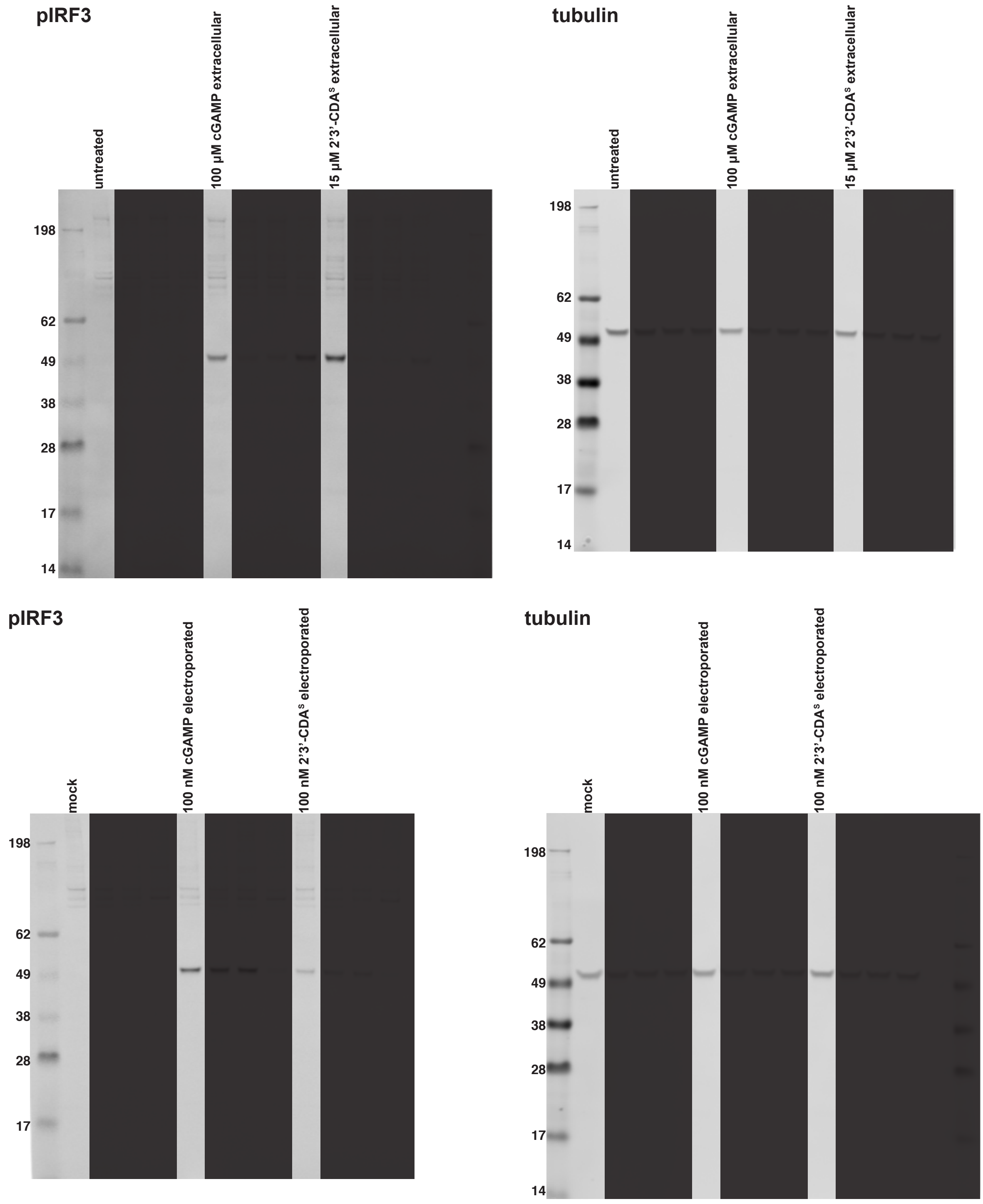


**Extended Supplementary Figure 5d -** Uncropped Western blots for Supplementary Figure 5d. Lanes not relevant to experiment are shaded.
